# Insights into the origin of metazoan multicellularity from predatory unicellular relatives of animals

**DOI:** 10.1101/817874

**Authors:** Denis V. Tikhonenkov, Elisabeth Hehenberger, Anton S. Esaulov, Olga I. Belyakova, Yuri A. Mazei, Alexander P. Mylnikov, Patrick J. Keeling

## Abstract

The diversity and biology of unicellular relatives of animals has strongly informed our understanding of the transition from single-celled organisms to the multicellular Metazoa. Here we analyse the cellular structures and complex life cycles of the novel unicellular holozoans *Pigoraptor* and *Syssomonas* (Opisthokonta). Both lineages are characterized by complex life cycles with a variety of cell types, the formation of multicellular aggregations and syncytium-like structures, and an unusual diet for single-celled opisthokonts (partial cell fusion and joint sucking of a large eukaryotic prey), all of which provide new insights into the origin of multicellularity in Metazoa. The ability to feed on large eukaryotic prey could have been a powerful trigger in the formation and development both aggregative (e.g., joint feeding, which also implies signalling) and clonal (e.g., hypertrophic growth followed by palintomy) multicellular stages that played important roles in the emergence of multicellular animals.

## Introduction

The origin of animals (Metazoa) from their unicellular ancestors is one of the most important evolutionary transitions in the history of life. Questions about the mechanisms of this transformation arose about 200 years ago, but is still far from being resolved today. Most investigations on the origin of Metazoa have focused on determining the nature of the shared, multicellular ancestor of all contemporary animals (Moroz et al., 2014; Srivastava et al., 2008; 2010). However, even the branching order of early, non-bilaterian lineages of animals on phylogenetic trees is still debated: some consider either sponges (Porifera) (Feuda et al., 2017; Philippe et al., 2009; Simion et al., 2017;) or Ctenophora (Dunn et al., 2008; Ryan et al., 2013; Whelan et al., 2015) or Placozoa (Schierwater et al., 2009; Signorovitch et al., 2007;) to be the first branch of extant metazoans. While molecular clock-based studies and paleontological evidence indicate that multicellular animals arose more than 600 million years ago (Maloof et al., 2010; Sharpe et al., 2015), we know less about how animals arose. To establish the sequence of events in the origin of animals from unicellular ancestors, we also need to investigate their closest relatives, the unicellular opisthokont protists. Information on the diversity and biology of the unicellular relatives of animals, their placement within the phylogenetic tree of opisthokonts, and the identification of molecular and morphological traits thought to be specific for animals within their unicellular sisters, have all strongly informed our understanding of the transition from single-celled organisms to the multicellular Metazoa (King et al., 2008; Suga et al., 2013; Suga, Ruiz-Trillo, 2013; Torruella et al., 2015).

Until recently, only three unicellular lineages, the choanoflagellates, filastereans, ichthyosporeans, as well as *Corallochytrium limacisporum*, a mysterious marine osmotrophic protist described in association with corals, have been described as collectively being sisters to animals. Together with animals they form the Holozoa within the Opisthokonta (Aleshin et al., 2007; Lang et al., 2002; Torruella et al., 2015). These unicellular organisms have extremely variable morphology and biology. Choanoflagellates represent a species-rich group of filter-feeding, bacterivorous, colony-forming protists, which possess a single flagellum surrounded by a collar of tentacles (microvilli). They are subdivided into two main groups – the predominantly marine Acanthoecida and the freshwater and marine Craspedida (Carr et al., 2017). Filastereans are amoeboid protists producing pseudopodia. Until recently, they were represented by only two species: the endosymbiont of a freshwater snail, *Capsaspora owczarzaki*, and the free-living marine heterotroph, *Ministeria vibrans* (Hertel et al., 2002; Shalchian-Tabrizi et al., 2008), which was recently shown to also possess a single, real flagellum (Mylnikov et al., 2019). Ichthyosporeans are parasites or endocommensals of vertebrates and invertebrates characterized by a complex life cycle, reproduction through multinucleated coenocytes colonies and flagellated and amoeboid dispersal stages (Suga, Ruiz-Trillo, 2013). *Corallochytrium* is a unicellular coccoid organism, which produces rough, raised colonies and amoeboid limax-like spores (Raghukumar, 1987.). Additionally, molecular data predict a cryptic flagellated stage for *Corallochytrium* (Torruella et al., 2015).

A large number of hypotheses about the origin of multicellular animals have been proposed. The most developed model for the origin of metazoan multicellularity is based on a common ancestor with choanoflagellates (James-Clark, 1866; Ivanov, 1967; King et al., 2008, Mikhailov et al., 2009; Nielsen, 1987, Ratcliff et al., 2012). This idea was initially based on the observed similarity between choanoflagellates and specialized choanocyte cells in sponges. Molecular investigations also supported the idea by consistently indicating that choanoflagellates are the closest sister group to Metazoa. However, molecular phylogeny itself does not reveal the nature of ancestral states, it only provides a scaffolding on which they might be inferred from other data. The evolutionary positions of the other unicellular holozoans (filastereans, ichthyosporeans, and *Corallochytrium*) are less clear and sometimes controversial (e.g. Cavalier-Smith, Chao, 2003; del Campo, Ruiz-Trillo, 2013; Hehenberger et al., 2017; Medina et al., 2003; Ruiz-Trillo et al., 2008; Shalchian-Tabrizi et al., 2008; Torruella et al., 2012, 2015;).

As noted above, many molecular traits that were thought to be “animal-specific” are now known to be present in unicellular holozoans, while conversely the loss of other traits have been shown to correlate with the origin of the animals. But gene content alone is not sufficient to provide a comprehensive understanding of the cell biology, life cycle and regulation capabilities of the unicellular ancestor, it requires also analysis of the biology of the extant unicellular relatives of animals (Sebé-Pedrós et al., 2017).

Recently we described a phylogenomic and transciptome analyses of three novel unicellular holozoans (Hehenberger et al., 2017), which are very similar in morphology and life style but not closely related. *Pigoraptor vietnamica* and *Pigoraptor chileana* are distantly related to filastereans, and *Syssomonas multiformis* forms a new phylogenetic clade, “Pluriformea”, with *Corallochytrium*. Both new genera of unicellular holozoans form the shortest and most slowly evolving branches on the tree, which improved support for many nodes in the phylogeny of unicellular holozoans. Also, comparison of gene content of the new taxa with the known unicellular holozoans revealed several new and interesting distribution patterns for genes related to multicellularity and adhesion (Hehenberger et al., 2017).

Here we report the detailed morphological and ultrastructural analyses of these new species, as well as describing their life cycle in culture, which are important implications for understanding the origin of animals as are the genetic analyses. All three species are shown to be predatory flagellates that feed on large eukaryotic prey, which is very unusual for unicellular Holozoa. They also appear to exhibit complex life histories with several distinct stages, including interesting multicellular structures that might offer important clues as to precursors of multicellularity. On the basis of these findings we discuss the current hypotheses about the origin of multicellular animals from their unicellular ancestors.

## Results and discussion

Detailed morphological descriptions of the cells and their aggregates are presented below. *Syssomonas multiformis* Tikhonenkov, Hehenberger, Mylnikov et Keeling 2017

### Morphology and life cycle

The organism is characterized by a large variety of life forms including flagellates, amoeboflagellates, amoeboid non-flagellar cells, and spherical cysts. The most common stage in the life cycle, a swimming flagellate cell, resembles a typical opisthokont cell, reminiscent of sperm cells of most animals and zoospores of the chytrid fungi. Cells are round-to-oval and propel themselves with a single, long posterior flagellum (Fig. 1A-C, X). The flagellum is smooth and emerges from the middle-lateral point of the cell, turns back and always directs backwardduring swimming. The cell rotates during swimming (Video 1). Flagellar beating can be very fast, which can create the appearance of two flagella. Motile flagellates can suddenly stop and change the direction of movement. The flagellated cells measure 7-14 µm in diameter. The flagellum length is 10-24, rarely 35 µm. Cyst diameter is 5 µm (Fig. 1D, Y).

**Fig.1.**
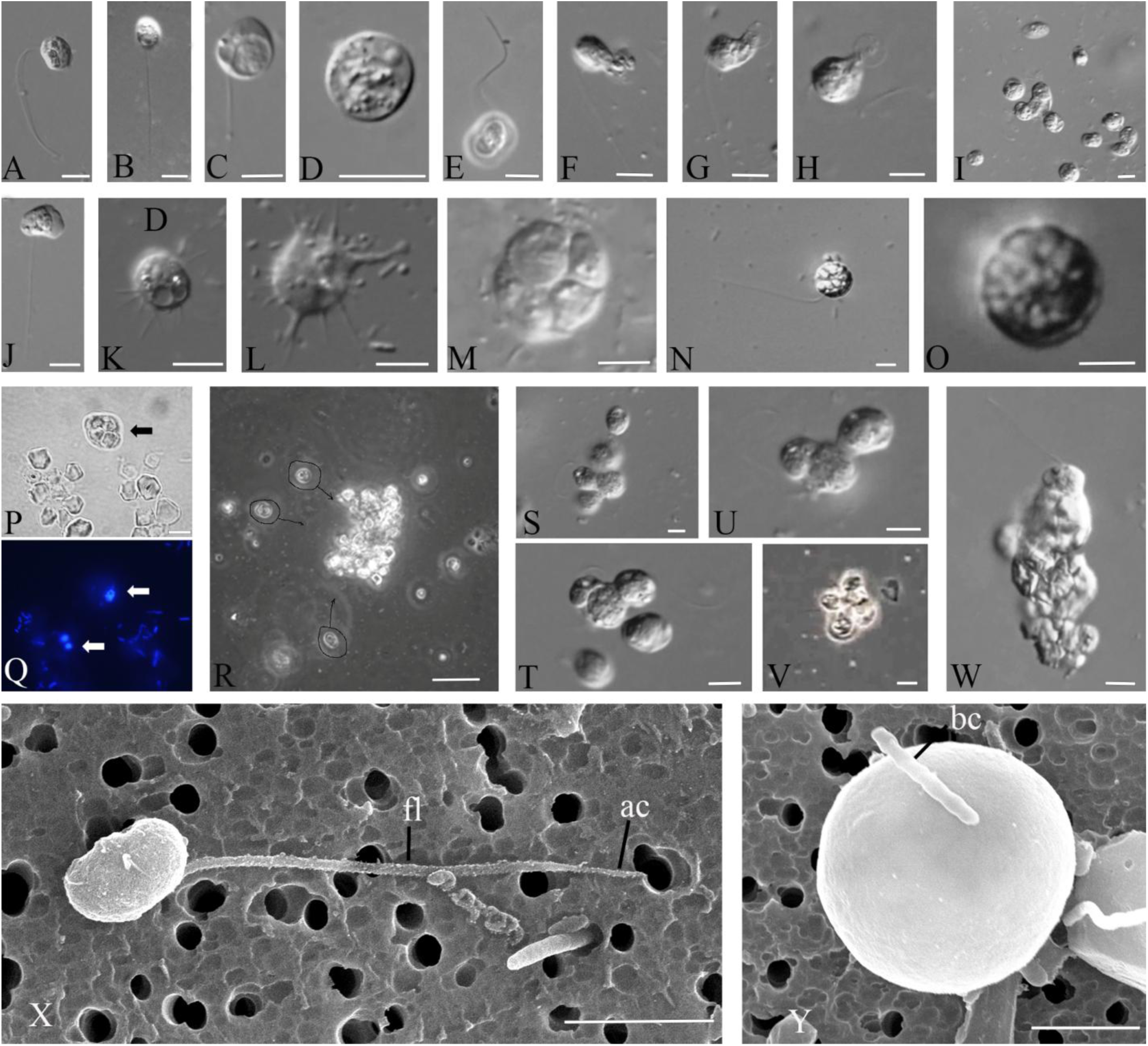
External morphology and life forms of *Syssomonas multiformis.* A-C – swimming flagellated cells; D – cyst; E – attached flagellated cell; F-H – sucking of eukaryotic prey; I – simultaneous joint feeding of three cells of *Syssomonas* on one prey cell with attaction of other speciment to the feeding spot; J – amoeboflagellate; K,L – amoeboid cell; M – palintomic cell-division inside the cyst; N – cell with inside vesicules; O – cyst with vesicules; P,Q – cells of *Syssomonas* engulfed starch granules (bright field (P) and fluorescent microscopy, DAPI staining); R – cells of *Syssomonas* with engulfed starch granules hiding into the starch druse, S-U, W – cell aggregations of *Syssomonas* near the bottom of Petri dish; V – floating aggregation of flagellated cells; X – general view of flagellated cell (SEM), Y – cyst (SEM). Scale bars: A-P, S-W – 10 μm, R – 100 μm, X – 3 μm, Y – 2 μm.

Solitary cells of *Syssomonas* can temporary attach to the substrate by the anterior part of the cell body. They produce water flow by rapid flagellum beating posteriorly and in that state resemble cells of choanoflagellates or choanocytes from sponges (Fig. 1E, Video 2).Floating flagellated cells can also move to the bottom and transform to amoeboflagellates (Fig. 1J, Video 3) by producing both wide lobopodia and thin short filopodia. Flagellar beating becomes slower and then stops. Amoeboflagellates crawl along the surface using their anterior lobopodia and can take up clusters of bacteria. The organism can lose the flagellum via three different modes: the flagellum may be abruptly discarded from its proximal part of the cell; a stretched flagellum may be retracted into the cell; the flagellum may convolve under the cell-body and then retract into the cell as a spiral (Video 4). As a result *Syssomonas* turns into an amoeba (Fig. 1K,L, Video 4). Amoeboid cells produce thin, relatively short filopodia and sometimes have two contractile vacuoles. Amoeboid cells are weakly motile. The transformation of amoeboflagellates and amoebae back to flagellates was also observed.

Amoeboid cells can also retract their filopodia, become roundish and transform into a cyst (Fig. 1D, Video 5). Palintomic divisions may occur inside the cyst and up to 16 (2, 4, 8, or 16) flagellated cells are released as a result (Fig. 1M, Video 6). Division into two cell structures was also observed in culture (Video 7), but it is hard to tell whether a simple binary longitudinal division of a *Syssomonas* cell with retracted flagellum has taken place, or the final stage of a division inside the cyst has been observed.

Floating, flagellated cells containing vesicular structures were observed (Fig. 1N, Video 8), however the process of formation and the purpose of these vesicles is unknown. After some time such cells lose their flagellum and transform into vesicular cysts with a thick cover (Fig. 1O). Division inside vesicular cysts was not observed within 10 days of observation. Such structures could represent resting cysts or dying cells containing autophagic vacuoles (the partial destruction of one such cyst was observed after 4 days of observation, see Video 8).

The organism is a predator; it takes up other flagellates (e.g. *Parabodo caudatus* and *Spumella* sp.) which can be smaller, about the same size, or larger than *Syssomonas.* But in contrast to many other eukaryotrophic protists, *Syssomonas* does not possess any extrusive organelles for prey hunting. After initial contact, *Syssomonas* attaches to the prey cell and sucks out their cytoplasm (without ingesting the cell membrane) (Fig. 1F-H, Video 9). The organism feeds better on inactive, slow moving or dead cells and can also capture intact prey cells and cysts by means unobserved. After attaching to the prey, many other *Syssomonas* cells become attracted to the same prey cell (likely by chemical signaling) and try to attach to it. Joint feeding was observed: several cells of *Syssomonas* can suck out the cytoplasm of the same prey cell together (Fig. 1I, Video 9).

In culture, *Syssomonas* can take up starch granules from rice grains, the granules can be the same size as the cells (Fig. 1P,Q). In the presence of *Syssomonas* cells, rice grains in Petri dishes crumble into small fragments and separate granules of starch (Fig. S1). Cells of *Syssomonas* with engulfed starch granules can hide within the starch druses and lose the flagellum (Fig. 1R). Numerous cysts integrated into the starch matrix were often observed in culture.

**Fig. S1.**
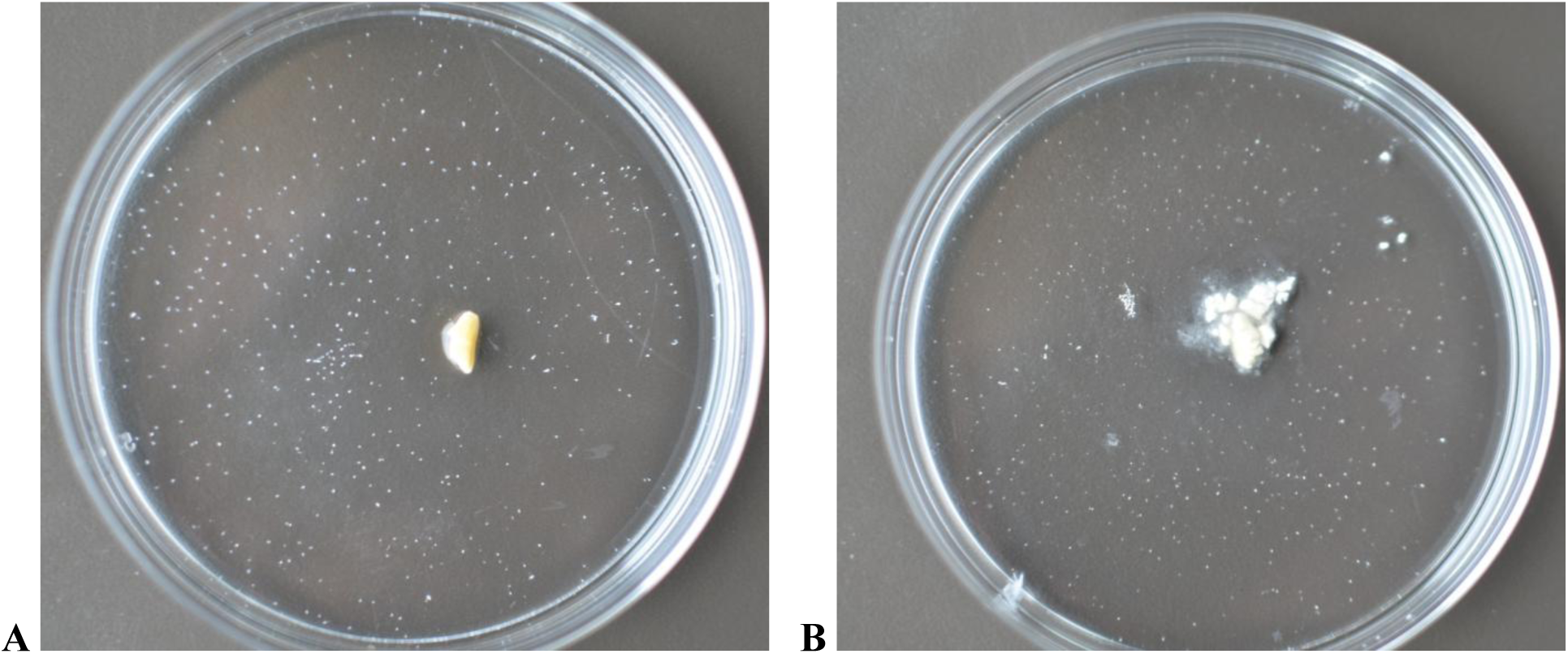
Rice grain destruction in Petri dish with Pratt medium and presence of the cells of *Parabodo caudatus* (prey) only (A) and *Syssomonas multiformis* (B) after 9 days of incubation.

The organism can also feed on clusters of bacteria (Video 10) using short pseudopodia. After feeding, *Syssomonas* cells become 2-3 times bigger and a large food vacuole is formed at the posterior end of the cell-body (Fig. 1C). In the absence of eukaryotic prey (cultivation on bacteria and/or rice grain/starch only), *Syssomonas* either dies or forms resting cysts. Bacteria alone are not sufficient food for *Syssomonas.*

Solitary cells of *Syssomonas* can partially merge and form temporary cell aggregations. They are usually shapeless, observed near the bottom, and consist of about 3-10 flagellated or non-flagellated cells (Fig. 1S-U, Video 11). Another type of aggregation is formed by only flagellated cells with outwards-directed flagella that can float in the water column and resemble the rosette-like colonies of choanoflagellates (Fig. 1V, Video 12). Both types of aggregations break up easily and it seems that the membranes of such aggregated cells are not fused.

However, in rich culture, solitary cells of *Syssomonas* can sometimes merge completely at the bottom of the Petri dish and form syncytium-like (or pseudoplasmodium) structures (it seems that nuclei do not merge after cell fusion). The budding of young flagellated daughter cells from such syncytia was observed (Video 13).Such syncytial structures with budding daughter cells have not been observed in other eukaryotes, to our knowledge, but multinucleated structures arising as a result of multiple cell aggregations or fusions of uninuclear cells are also known in Dictyostelia (Eumycetozoa) and *Copromyxa* (Tubulinea) in the Amoebozoa (sister group of Opisthokonta), as well as in other protists, such as Acrasidae in the Excavata, *Sorogena* in the Alveolata, *Sorodiplophrys* in the Stramenopiles, and *Guttulinopsis* in the Rhizaria (Brown et al., 2012). Within the opisthokonts, aggregation of amoeboid cells is only known in the sorocarpic species *Fonticula alba* (Holomycota) (Brown et al., 2009). We should also note that a syncytium is not an unusual cell structure in many fungi and animals; e.g. most of the cytoplasm of glass sponges (Hexactinellida), the teguments of flatwormsas well as the skeletal muscles and the placenta of mammals (Gobert et al., 2003; Leyset al., 2006) have a syncytial structure.

In *Syssomonas*, the processes of cells merging attracts (again, likely by chemical signalling) many other cells of *Syssomonas*, which actively swim near aggregates or syncytium-like structures and try to attach to them. Some of the these cells succeed to merge and the aggregates grow.

All aggregations and syncytial-like structures do not seem to form by cell division, but rather by a merger of the population of cells in the culture (although all cells in the clonal culture are offspring of a single cell of *Syssomonas*).

All of the above-described life forms and cellular changes do not represent well-defined phases of the life cycle of *Syssomonas*, but rather embody temporary transitions of cells in culture which are reversible.

*Syssomonas* grows at room temperature (22°C) and can survive at temperatures from +5 to 36 °C. At high temperature (30-35 °C) the prey cells (bodonids) in culture become immobile and roundish; *Syssomonas* actively feeds on such easily accessible cells, multiplies and produces high biomass. In the absence of live eukaryotic prey, increasing the incubation temperature does not lead to an increase in cell numbers. The cells grow at pH values from 6 to 11. Agitation of culture does not lead to the formation of cell aggregates as was observed in the filasterean *Capsaspora* (Sebé-Pedrós et al., 2013b).

### Cell ultrastructure

The cell is naked and surrounded by the plasmalemma. The naked flagellum ends in a short, narrowed tip – the acroneme (Fig. 1X, 2D). A single spiral or other additional elements (e.g. a central filament typical for choanoflagellates) in the transition zone of the flagellum were not observed (Fig. 2B, 2C). The flagellar axoneme has an ordinary structure (9+2) in section (not shown). The flagellum can be retracted into the cell (Fig. 2E). A cone-shaped rise at the cell surface around the flagellum base was observed (Fig. 2B, 2C). The flagellar transition zone contains a transversal plate which is located above the cell surface (Fig. 2B).

**Fig. 2.**
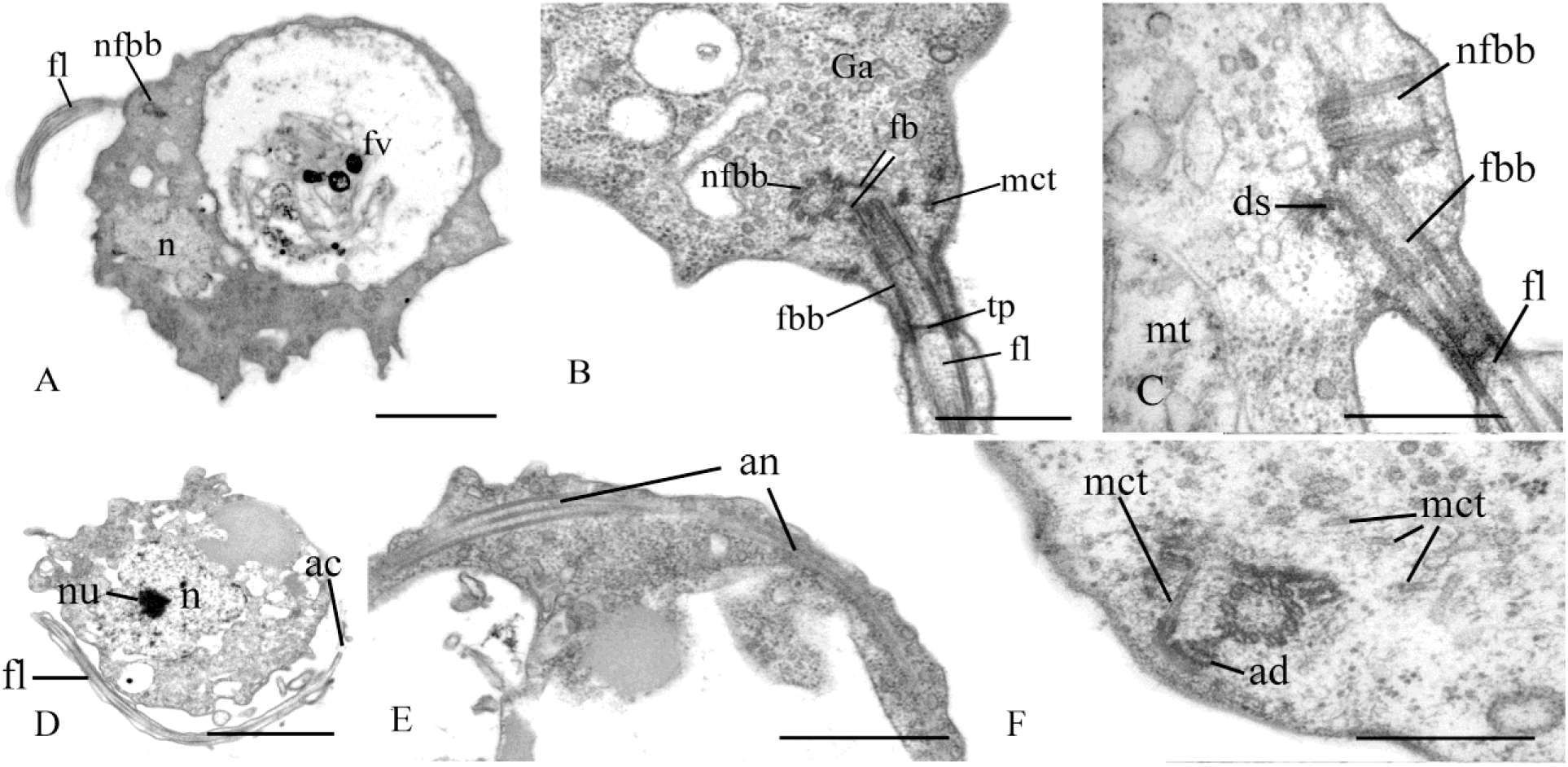
General view and flagellum structure of *Syssomonas multiformis* (TEM). A – general view of the cell section, B,C – arrangement of flagellum and basal bodies, D – twisting of the flagellum around the cell, E – retracted flagellum axoneme inside the cell, F – basal body of the flagellum and nearest structures. ac – acroneme, ad – arc-like dense structure, an – flagellum axoneme, ds – dense spot, fb – fibril, fbb – flagellar basal body, fl – flagellum, fv – food vacuole, Ga – Golgi apparatus, mct – microtubule, mt – mitochondrion, n – nucleus, nfbb – non-flagellar basal body, nu – nucleolus, tp – transversal plate. Scale bars: A – 2, B – 0.5, C – 0.5, D – 2, E – J – 0.5 μm.

Two basal bodies, one flagellar and one non-flagellar (Fig. 2A–C), lie approximately at a 45-90 degrees angle to each other (Fig. 2B, 2C). The flagellar root system consists of several elements. Arc-like dense material, representing satellites of the kinetosome, is connected with the flagellar basal body and initiates microtubules which run into the cell (Fig. 2F). Radial fibrils originate from the flagellar basal body (Fig. 3A-C, 3G) and resemble transitional fibres. At least two fibrils connect to the basal bodies (Fig. 2B). It can be seen from serial sections that microtubules originate near both basal bodies (Fig. 3 A–F). Dense (osmiophilic) spots are situated near the basal bodies and some of them initiate bundles of microtubules (Fig. 3 I,J,L). Microtubules originating from both basal bodies singly or in the form of a fan probably run into the cell (Fig. 2 B, Fig. 3F–K). One group of contiguous microtubules begins from the dense spot (Fig. 3L) and goes superficially close to the plasmalemma (Fig. 3 E,F,L).

**Fig. 3.**
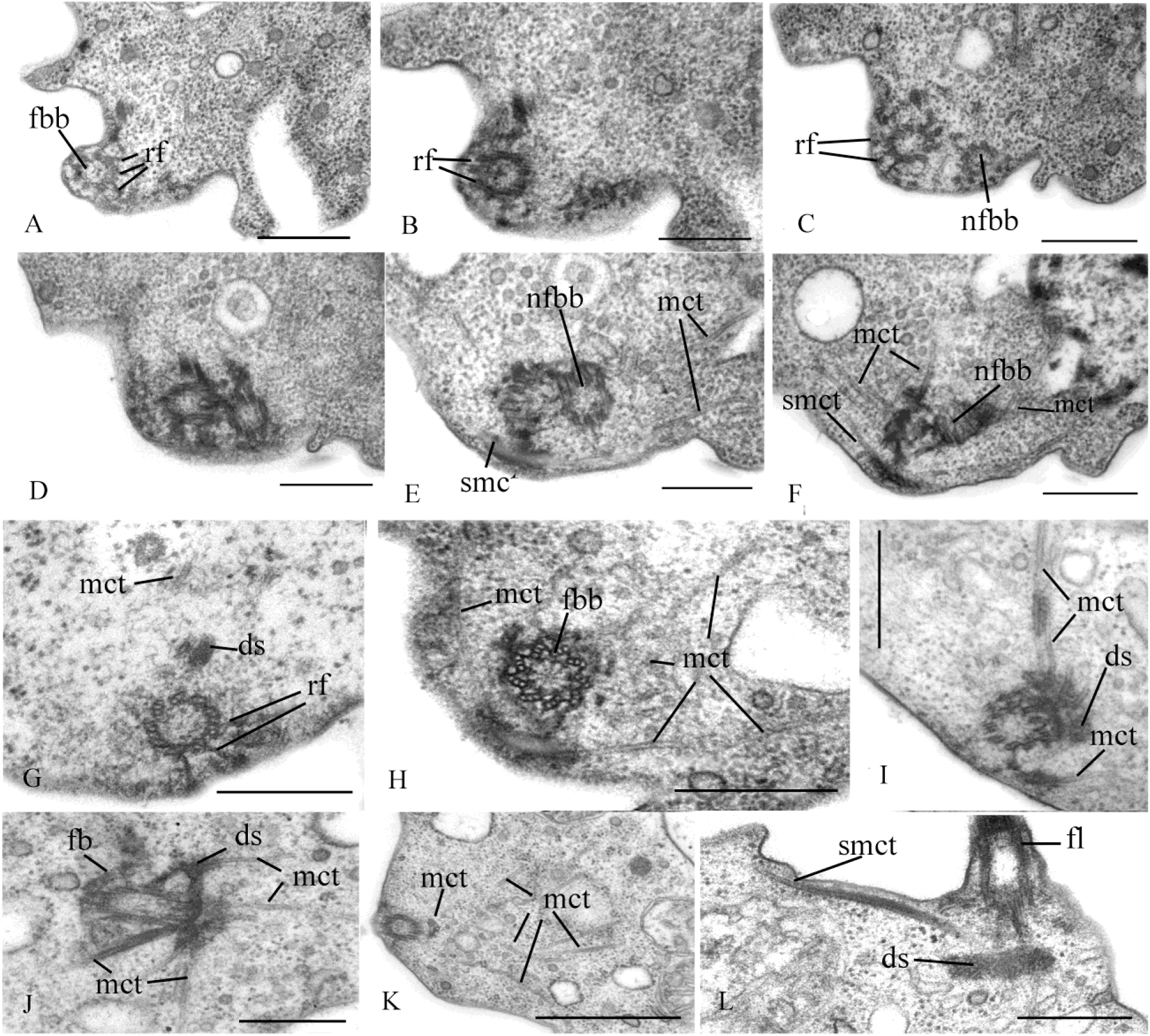
Arrangement of kinetosomes of *Syssomonas multiformis.* A – F – serial sections of the kinetosomal area, G–L – structures nearby the kinetosomes. ds – dense spot, fb – fibril, fbb – flagellar basal body, mct – microtubule, nfbb – non-flagellar basal body, rf – radial fibrils, smmt – submembrane microtubules. Scale bars: A–J, L – 0.5, K – 1 μm.

The nucleus is 2.6 µm in diameter, has a central nucleolus and is situated closer to the posterior part of the cell (Fig. 2 A, D, Fig. 4 H). The Golgi apparatus is of usual structure and is positioned close to the nucleus (Fig. 2B, Fig. 4A). The cell contains several mitochondria with lamellar cristae (Fig. 4 B–D). Unusual reticulate or tubular crystal-like structures of unknown nature were observed inside the mitochondria (Fig. 4C, D). A contractile vacuole is situated at the periphery of the cell and is usually surrounded by small vacuoles (Fig. 4E).

**Fig. 4.**
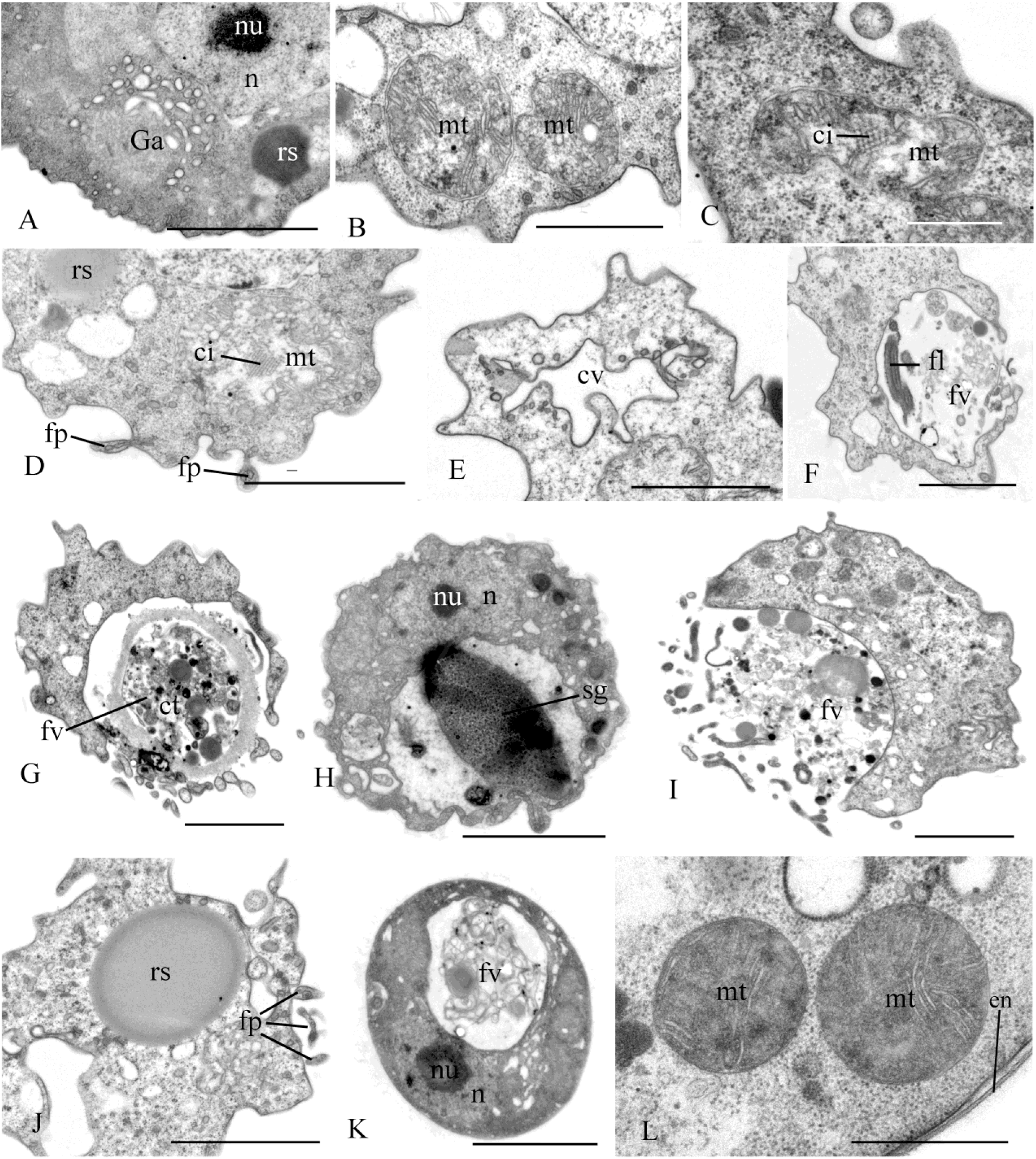
Cell structures and organelles of *Syssomonas multiformis.* A – Golgi apparatus, B – D – mitochondria, E – contractile vacuole, F – food vacuole with remnants of eukaryotic prey (*Parabodo*), flagella and paraxial rods are seen, G – food vacuole containing cyst of *Parabodo*, H – food vacuole containing starch granule, I – Exocytosis of food vacuole, J – reserve substance and filopodia, K – L – cysts. ci – crystalloid inclusion, ct – cyst of the prey cell, cv – contractile vacuole, en – cyst envelope, fl – flagellum, fp – filopodia, fv – food vacuole, ga – Golgi apparatus, mt – mitochondrion, n – nucleus, nu – nucleolus, rs – reserve substance, sg – starch granule. Scale bars: A – 2, B – 1, C – 0.5, D – 2, E – 2, F – 2, G – 2, H – 2, I – 2, J – 2, K – 2, L – 1 μm.

A large food vacuole is usually located posteriorly or close to the cell center and contains either remnants of eukaryotic prey, e.g. cells (paraxial flagellar rods are seen) or cysts (fibrous cyst envelope is seen) of *Parabodo caudatus*, or starch granules (Fig. 2A, Fig. 4F–H). Exocytosis occurs on the posterior cell end (Fig. 4I).

Thin filopodia are located on some parts of cell surface (Figs. 4D, 4J).

Storage compounds are represented by roundish (presumably glycolipid) granules 0.8 μm in diameter (Fig. 4 A,D,J).

A flagellum or flagellar axoneme, or two kinetosomes, as well as an eccentric nucleus, mitochondria with lamellate cristae and dense matrix, and a food vacuole with remnants of the prey cells are all visible inside cysts containing dense cytoplasm (Fig. 4K, L).

Extrusive organelles for prey hunting were not observed in any cell type.

*Pigoraptor vietnamica* Tikhonenkov, Hehenberger, Mylnikov et Keeling 2017

### Morphology and life cycle

The uniflagellated, elongated-oval cells are 5-12 µm long (Fig. 5 A,B,H,I). The flagellum length is 9-14 µm. Saturated cells with a large food vacuole become roundish. The body plan, movement, feeding, and growth conditions of *Pigoraptor* are identical to *Syssomonas multiformis*, except for the feeding on starch granules, which was not observed in *Pigoraptor*.

**Fig. 5.**
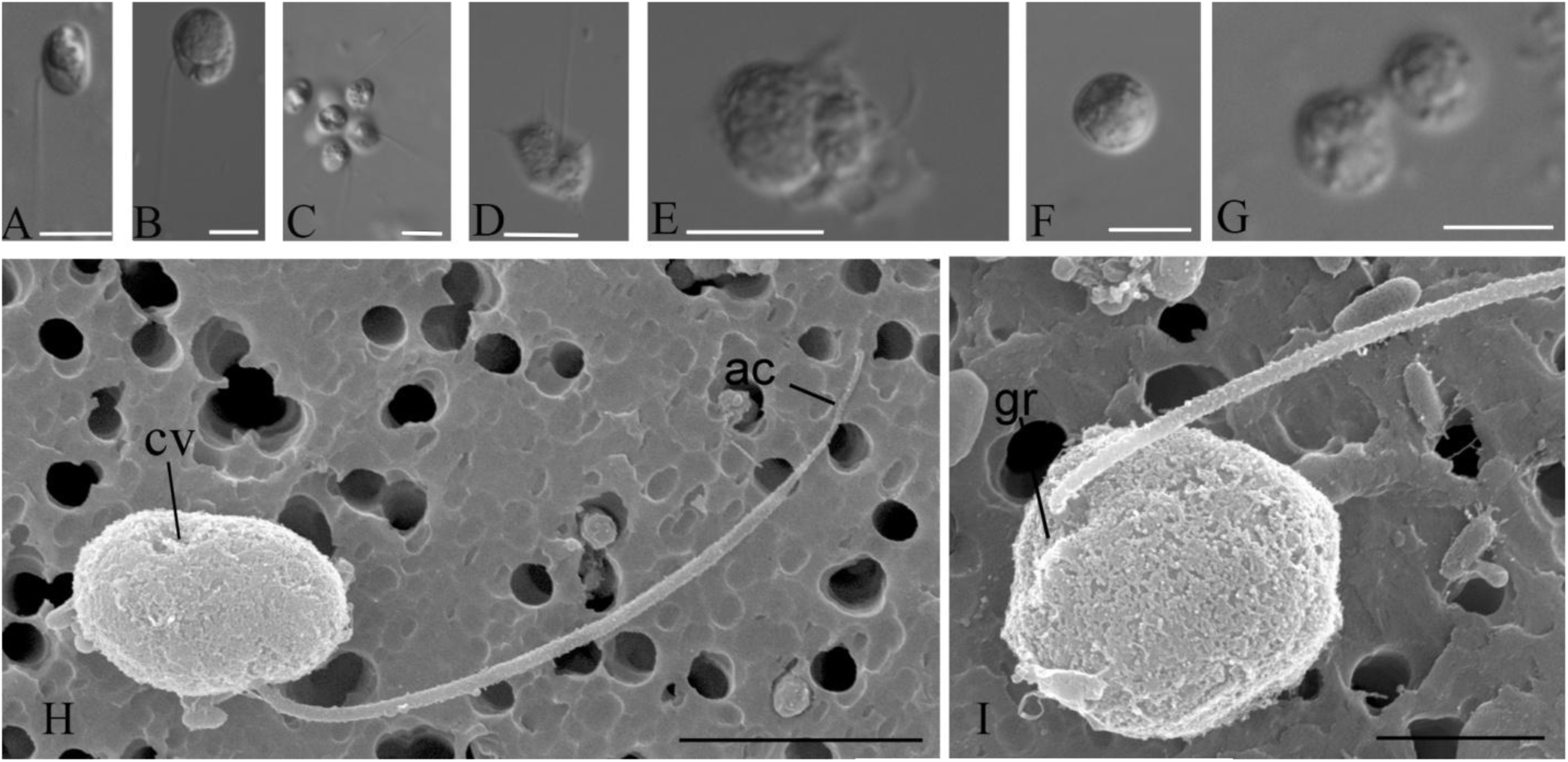
External morphology and life forms of *Pigoraptor vietnamica.* A, B, H, I – general view of the cell (DIC and CЭM); C – aggregation of flagellated cells; D – amoeboflagellate with filopodia; E – cell with lobodopia; F – cyst; G – binary division. ac – acroneme, cv – contractile vacuole, gr – groove. Scale bars: A-G – 10; H – 4, I – 2 μm.

The main stage of the life cycle is a swimming flagellated cell, which can form thin long, sometimes branching filopodia that can attach to the substrate (Fig. 5D). Wide lobopodia were also observed on some cells (Fig. 5E). Non-flagellate crawling amoebas were not observed. *Pigoraptor* cells can retract the flagellum and become roundish. After several hours, such spherical cells either divide into two daughter cells or turn into cysts (Fig. 5F), which stay intact for a long period. Binary division was observed also inside the cyst (Fig. 5G), resulting in two daughter cells that produce flagella and disperse.

Cells of *Pigoraptor vietnamica* also form easily disintegrating aggregations (Fig. 5C) and feed jointly (Video 14). The adjacent cells can partially merge during feeding. These processesalso seem to attract many other cells of *Pigoraptor.*

### Cell ultrastructure

A single, naked flagellum with an acroneme originates from a small lateral groove and directs backward (Fig. 5 H,I). The cell is naked and surrounded by the plasmalemma. Two basal bodies, flagellar and non-flagellar, are located near the nucleus, lie approximately at a 90 degrees angle to each other and are not connected by visible fibrils (Fig. 6 A,B; Fig. 7 A,B; Fig. 8 A–F). The flagellum axoneme has an ordinary structure (9+2) in section (Fig. 6 D,E).A thin central filament, which connects the central pair of microtubules to the transversal plate, was observed (Fig. 6 C,F).The flagellum can be retracted into the cell (Fig. 9C).The flagellar root system is reduced. Radial fibrils arise from the flagellar basal body (Fig. 7C). Microtubules pass near the flagellar basal body (Fig. 7B, Fig. 8 E,F). Serial sections show that the non-flagellar basal body does not initiate the formation of microtubules (Fig. 8 A,B)

**Fig. 6.**
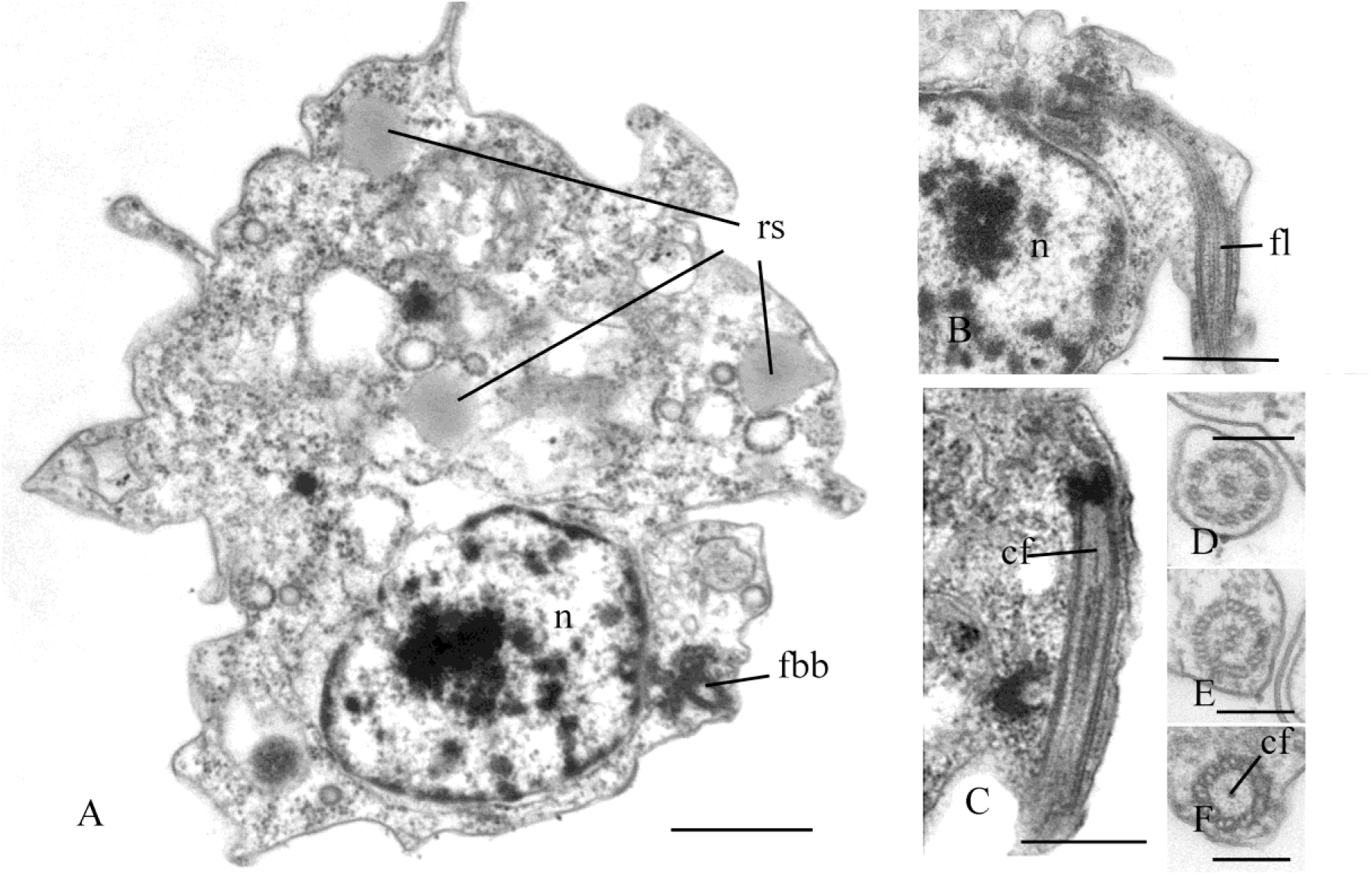
General view and flagellum structure of *Pigoraptor vietnamica*, TEM. A – longitudinal cell section. B–C – longitudinal section of flagellum, D–F – transverse flagellum sections in transitional area. cf – central filament, fbb – flagellar basal body, fl – flagellum, n – nucleus, rs – reserve substance. Scale bars: A – 1, B – 0.5, C – 0.5, D – F – 0.2 – μm.

**Fig. 7.**
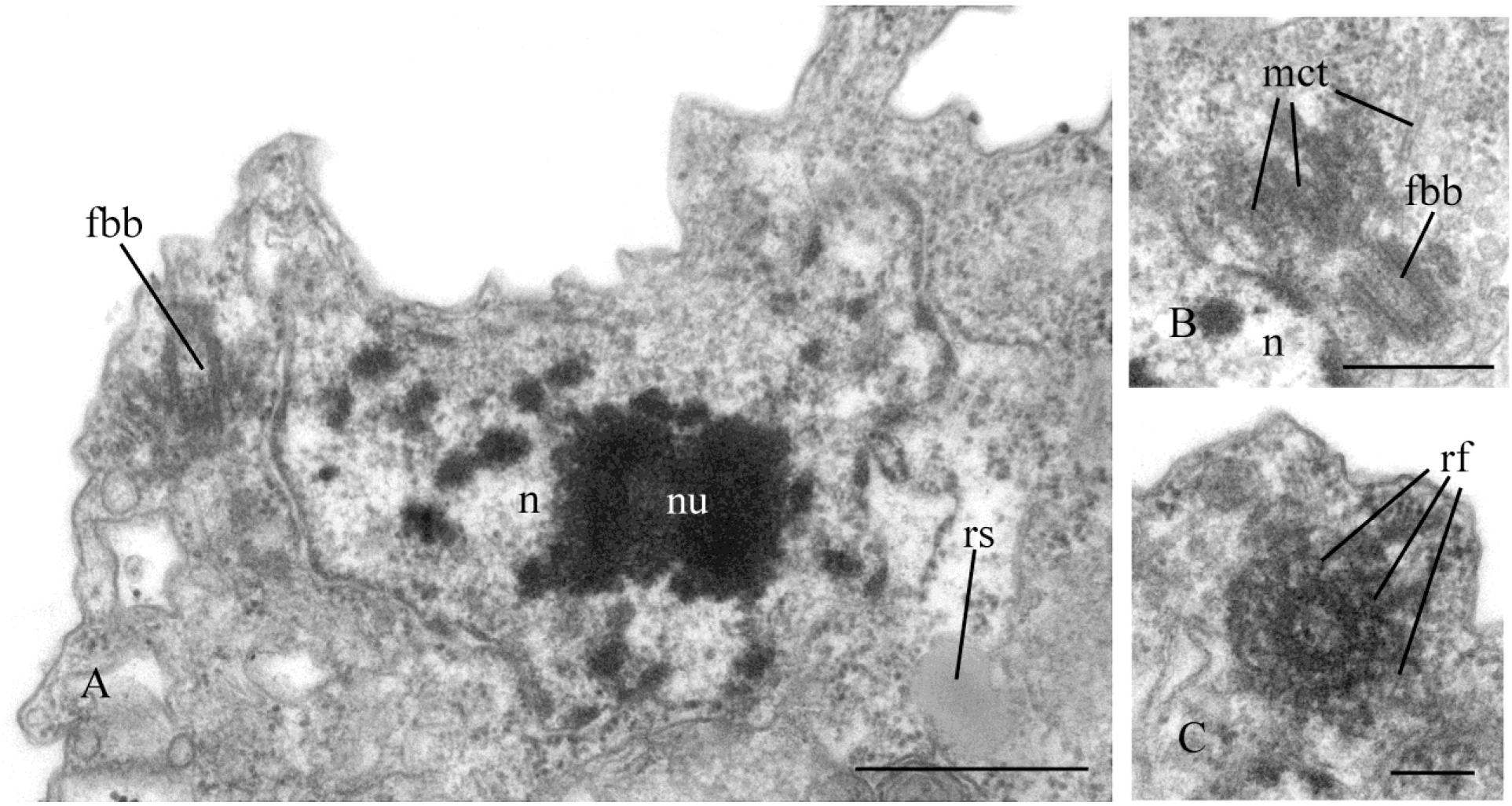
Nucleus and arrangement of basal bodies of *Pigoraptor vietnamica.* A – part of the cell containing nucleus and flagellar basal body, B – non-flagellar basal body, C – flagellar basal body and surrounding structures. fbb – flagellar basal body, mct – microtubule, n – nucleus, nu – nucleolus, rf – radial fibrils, rs – reserve substance. Scale bars: A – 0.5, B – 0.5, C – 0.2 μm.

**Fig. 8.**
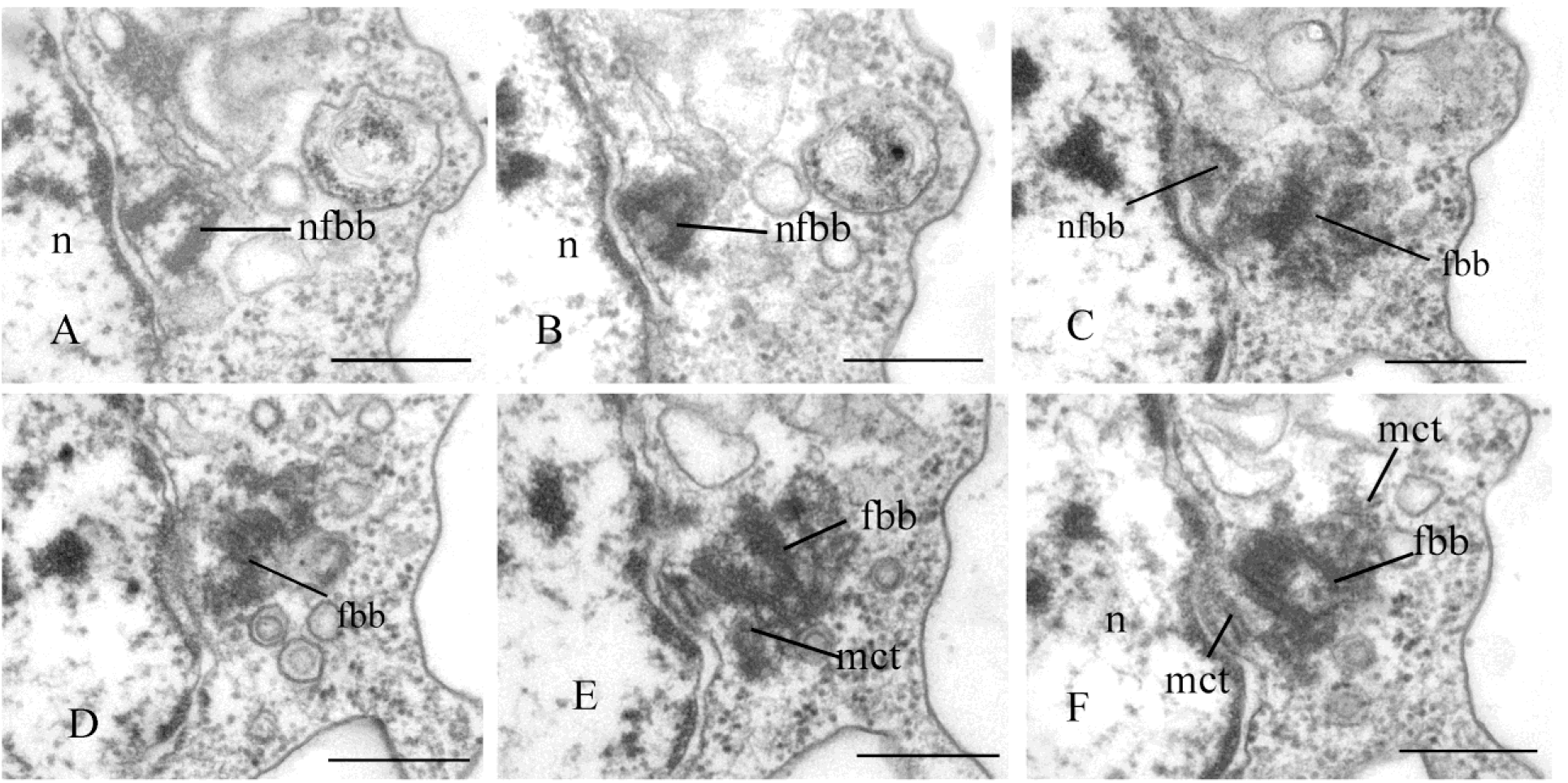
Arrangement of two basal bodies of *Pigoraptor vietnamica* relative to one another. A – F – serial sections of basal bodies. fbb – flagellar basal body, mct – microtubule, n – nucleus, nfbb – non-flagellar basal body. Scale bars: A – F – 0.5 μm.

**Fig. 9.**
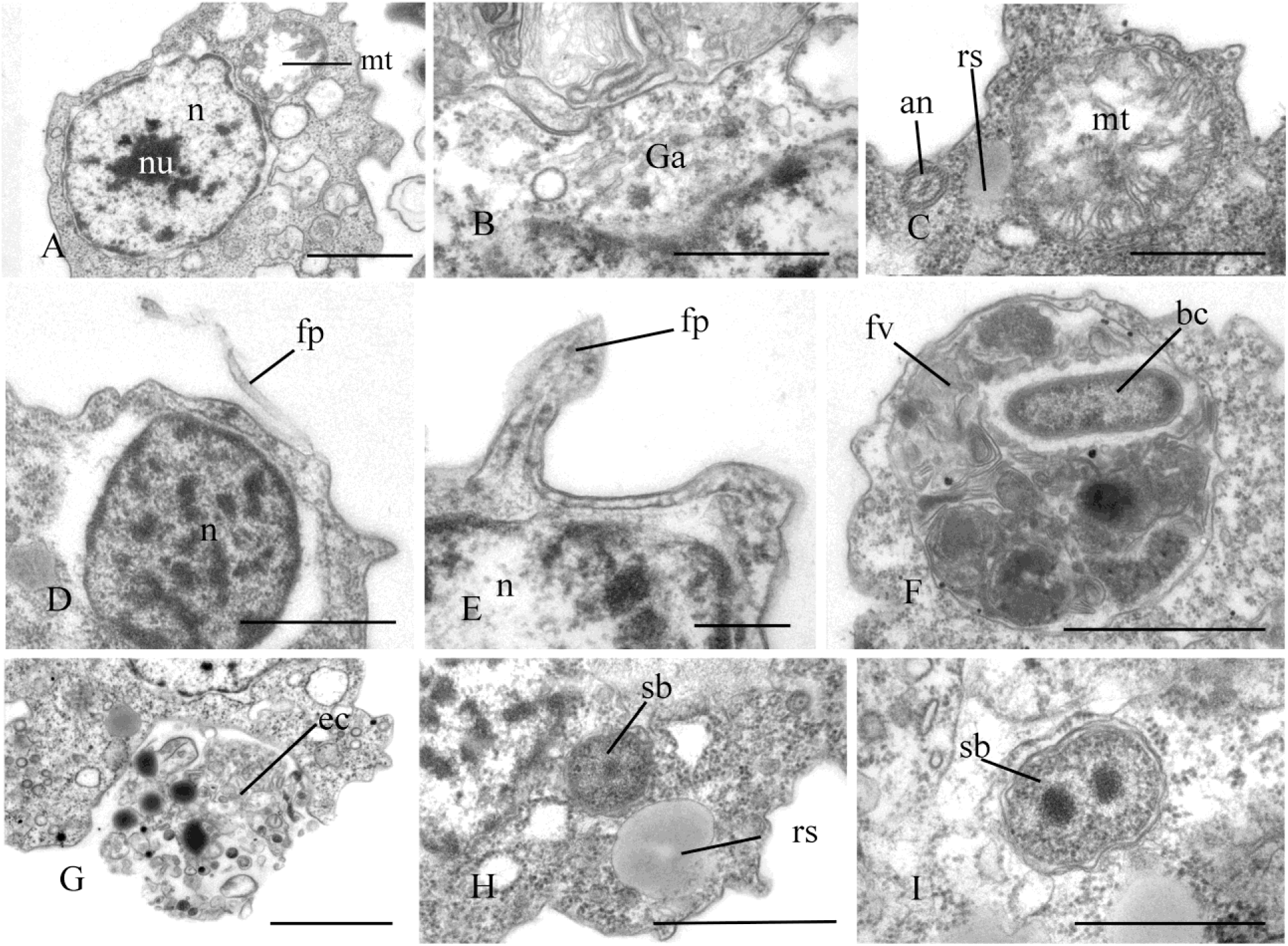
Sections of nucleus and other call structures of *Pigoraptor vietnamica.* A – nucleus, B – Golgi apparatus, C – mitochondrion, D – E – nucleus and filopodia, F – food vacuole, G – exocytosis, H – reserve substance, I – dividing symbiotic bacteria. an – flagellar axoneme, bc – bacterium, ec – ectoproct, fb – filopodium, fv – food vacuole, ga – Golgi apparatus, mt – mitochondrion, n – nucleus, nu – nucleolus, rs – reserve substance, sb – symbiotic bacteria. Scale bars: A – 1, B – 0.5, C – 0.5, D – 0.5, E – 0.2, F – 1, G – 1, H – 1, I – 0.5 μm.

The roundish nucleus is about 1.5 µm in diameter, contains a prominent nucleolus (Fig. 6 A,B, Fig. 7A, Fig. 9 A, D), and is situated closer to the posterior end of the cell. Chromatin granules (clumps) are scattered within the nucleoplasm. The Golgi apparatus is adjacent to the nucleus (Fig. 9B). Cells contain several mitochondria that possess lamellar cristae (Fig. 9 A,C). Rare thin filopodia have been observed on the cell surface (Fig. 9 D,E). Cells usually contain one large food vacuole (Fig. 9F), which contains remnants of eukaryotic prey and bacteria. Exocytosis takes place on the anterior cell end (Fig. 9G). Storage compounds are represented by roundish (presumably glycolipid) granules 0.3-0.4 μm in diameter (Fig. 6A, 7A, 9 C,H). Cells contain symbiotic bacteria, which are able to divide in the host cytoplasm (Fig. 9 H,I). A single contractile vacuole is situated close to the cell surface (not shown on cell sections but visualized by TEM, Fig. 5H).

*Pigoraptor chileana* Tikhonenkov, Hehenberger, Mylnikov et Keeling 2017

### Morphology and life cycle

The uniflagellated, roundish cells measure 6-14 µm in diameter. The flagellum emerges from a shallow groove and is 8-16 µm in length (Fig. 10 A,B,H,I). The flagellum ends with the acroneme. This species is identical to *Pigoraptor vietnamica* in body plan, movement, feeding, growth conditions, joint feeding and aggregation behaviours (Fig. 10 C, Video 15, 16), encystation (Fig. 10 D), binary division (Fig. 10 E), but additionally characterized by the absence of symbiotic bacteria and much reduced capability to produce filopodia and lobopodia (Fig. 10 F,G), which are extremely rare in *Pigoraptor chileana*.

**Fig. 10.**
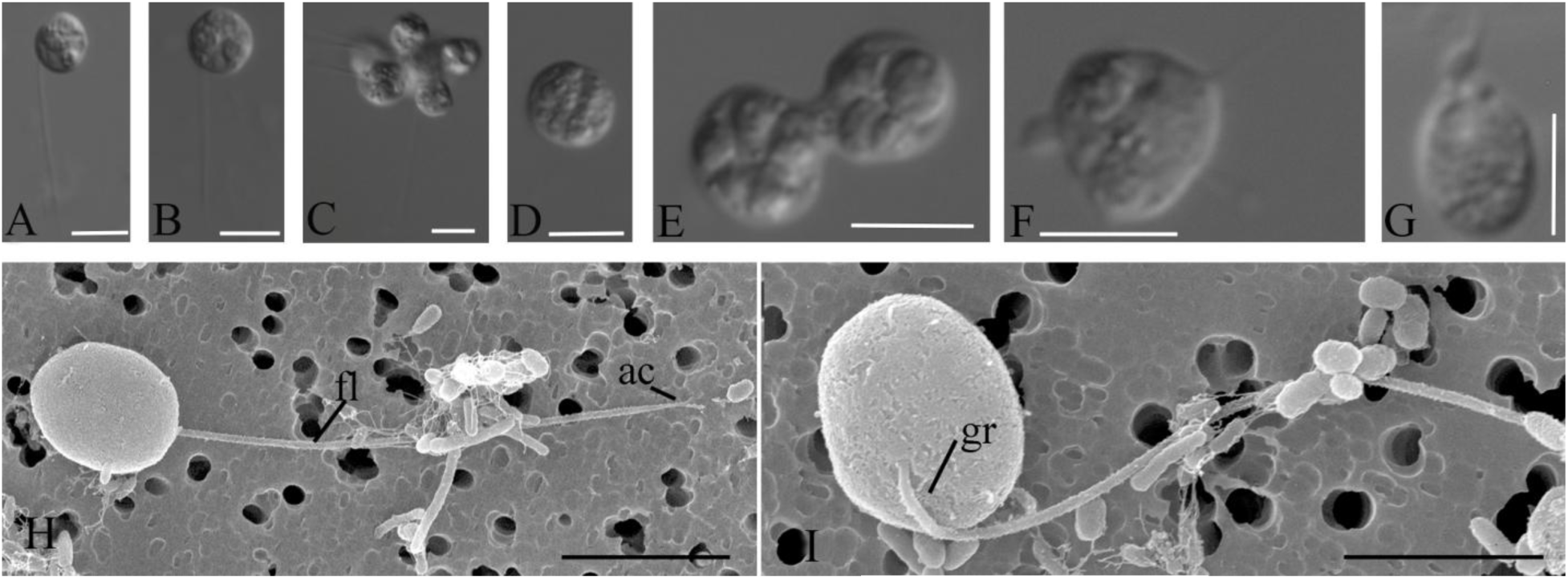
External morphology and life forms of *Pigoraptor chileana.* A, B, H, I – general view of flagellated cell (DIC and SEM); C – cell aggregation; D – cyst; E – binary division; F,G – cell with short lobopodia and filipodia. ac – flagella arconeme, gr – groove, fl – flagellum. Scale bars: A-G – 10, H – 9, I – 4 μm.

### Cell ultrastructure

The cell is naked and surrounded by the plasmalemma. The nucleus is positioned close to the posterior cell end (Fig. 11 A). The flagellum is naked and the flagellar axoneme has an ordinary structure (9+2) in section (Fig. 11 B–D). The flagellum can be retracted into the cell which is visible in some sections (Fig. 11 A, C). Flagellar and non-flagellar basal bodies are located near the nucleus (Fig. 11 A) and lie approximately at a 60-90 degrees angle to each other (Fig. 12 A–F).The flagellar basal body contains a wheel-shaped structure in the proximal part (Fig. 11 E,F). Single microtubules and microtubule bundles are situated near this basal body (Fig. 11 E–H). Some microtubules arise from dense spots close to the basal body (Fig. 11 H).

**Fig. 11.**
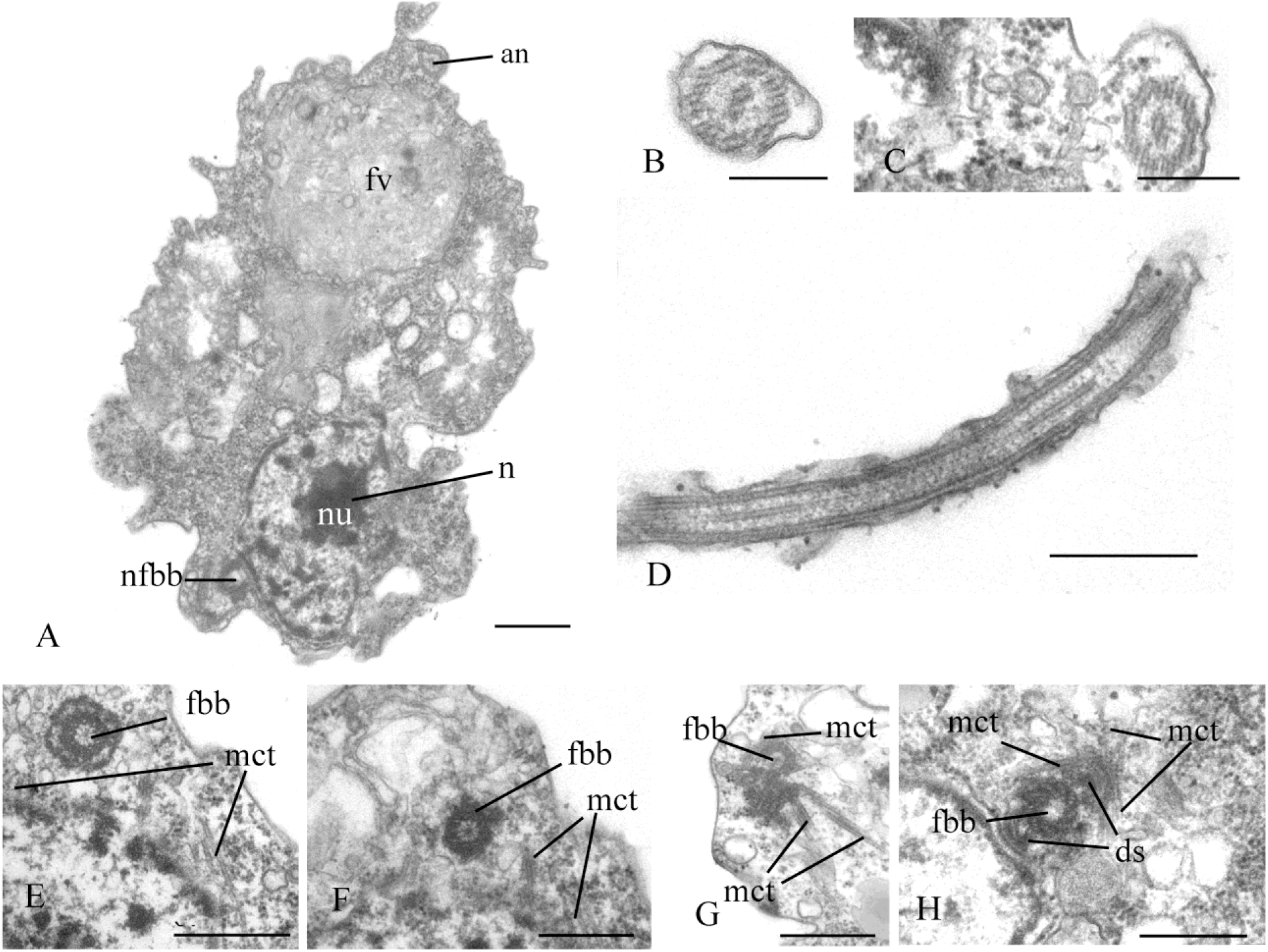
General view, flagellum and flagella root system of *Pigoraptor chileana.* A – general view of the cell section. B – D – flagellum, E– H – flagellar basal body and surrounding structures. an – axoneme, ds – dense spot, fbb – flagellar basal body, fv – food vacuole, mct – microtubule, n – nucleus, nfbb – non-flagellar basal body, nu – nucleolus. Scale bars: A – 0.5, B – 0.2, C – 0.2, D – 0.5, E – 0.5, F – 0.5, G – 0.5, H – 0.5 μm.

**Fig. 12.**
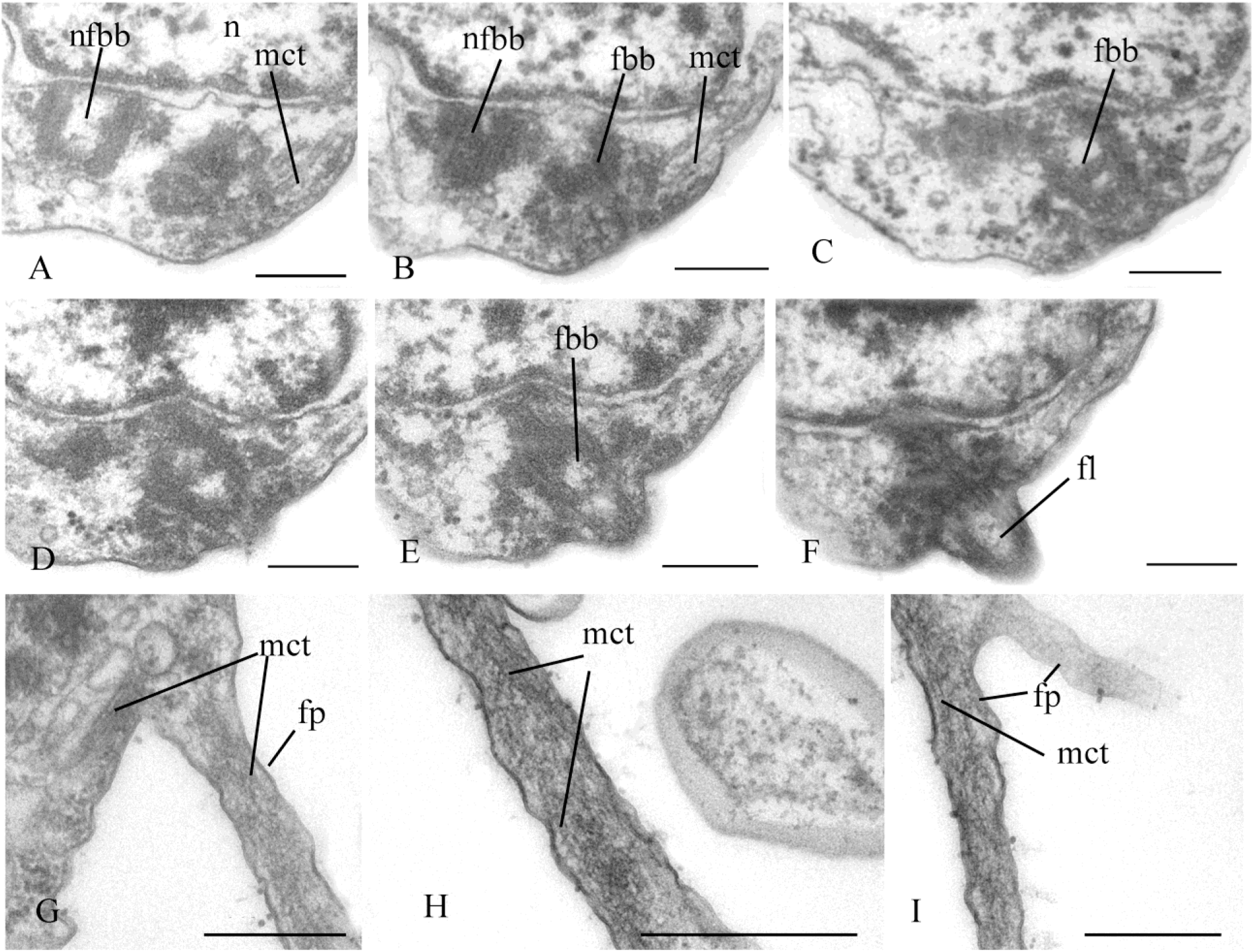
Arrangement of basal bodies and structure of filopodia of *Pigoraptor chileana.* A – F – serial sections of basal bodies, G – I – filipodia. fbb – flagellar basal body, fl – flagellum, fp – filopodium, mct – microtubule, n – nucleus, nfbb – non-flagellar basal body. Scale bars: A – F – 0.2, G – I – 0.5 μm.

Rare, thin, sometimes branching filopodia may contain superficially microtubule-like profiles (Fig. 12 G–I). The roundish nucleus is about 1.5 µm in diameter and has a central nucleolus (Fig. 11A). Chromatin granules are scattered within the nucleoplasm. The Golgi apparatus was not observed. Mitochondria contain lamellar cristae and empty space inside (Fig. 13 A, B). Cells usually contain one large food vacuole (Fig. 11A, Fig. 13 C), which contains remnants of eukaryotic prey and bacteria. Storage compounds are represented by roundish (presumably glycolipid) granules 0.2-0.4 μm in diameter (Fig. 13 C). The single ultrathin section of the dividing cell (possible open orthomitosis) was obtained in metaphase stage (Fig. 13 D).

**Fig. 13.**
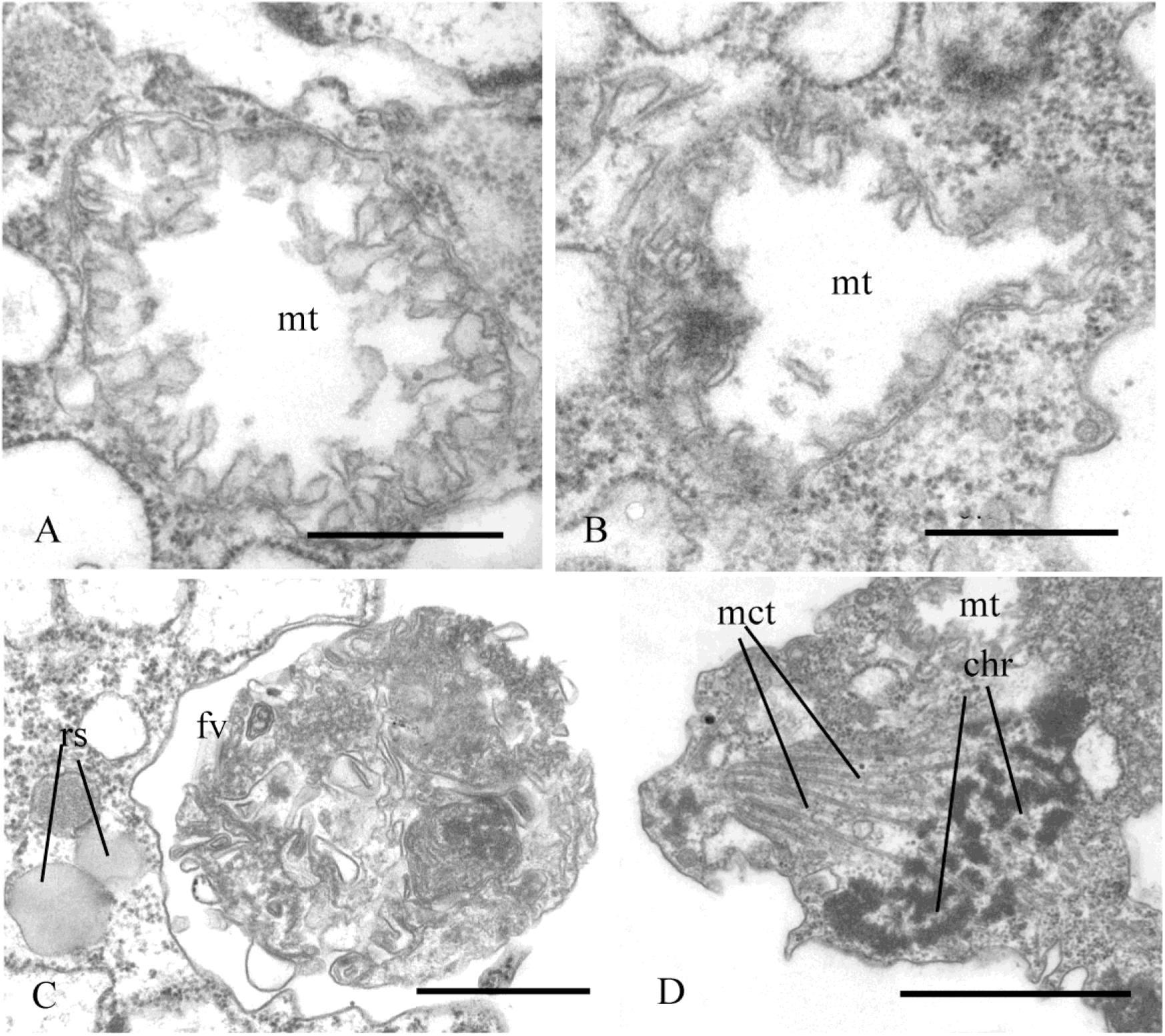
Mitochondria, food vacuole and nucleus division of *Pigoraptor chileana.* A, B – mitochondria, C – food vacuole and exocytose, D – nucleus division in metaphase stage. chr – chromosomes, fv – food vacuole, mct – microtubule, mt – mitochondrion, rs – reserve substance. Scale bars: A – 0.5, B – 0.5, C – 0.5, D – 1 μm.

### Key features of novel unicellular opisthokonts and origin of multicellularity in Metazoa

Our understanding of the origin and early evolution of animals has transformed as a result of the study of their most closely related sister groups of unicellular organisms: choanoflagellates, filastereans, and ichthyosporeans (King et al., 2008; Suga et al., 2013; Suga, Ruiz-Trillo, 2013). Our discovery of previously unknown unicellular Holozoa from freshwater bottom sediments in Vietnam and Chile provides new material for analysis.

#### Eukaryotrophy

A distinctive feature of all three new species is their feeding on eukaryotic prey of similar or larger size, which is unusual (if not unique) for unicellular Holozoa. While they consume entire prey cells or only the cytoplasmic contents of eukaryotic cells, which resembles the phagocytotic uptake of the contents of *Schistosoma mansoni* sporocysts by the filasterean

*Capsaspora owczarzaki* in laboratory conditions (Owczarzak et al., 1980), they can also feed on clusters of bacteria, resembling the phagocytic uptake of bacteria by choanoflagellates (Dayel, King, 2014). It is particularly noteworthy that the organisms we discovered, do not possess extrusive organelles for paralyzing and immobilizing the prey, which is typical for eukaryovorous protists. Our observations show that, prior to absorption, they somehow adhere to the surface of the prey cell. Studies on the choanoflagellate *Monosiga brevicollis* have shown that cadherins, that function as cell-cell adhesion proteins in animals, are located on the microvilli of the feeding collar and colocalize with the actin cytoskeleton (Abedin, King, 2008). *M. brevicollis* is non-colonial, thus suggesting that cadherins participate in prey capture, not colony formation. In addition, studies of the colonial choanoflagellate *Salpingoeca rosetta* did not indicate a role of the cadherins in colony formation, further supporting the notion that cadherins do not play a role in cell-cell adhesion between choanoflagellates, and perhaps also did not in the unicellular ancestor of animals (Sebé-Pedrós et al., 2017). In the case of the unicellular predators *Syssomonas* and *Pigoraptor*, adherence to a large and actively moving prey seems to be crucial for feeding and important for survival. Interestingly, the *Syssomonas* transcriptome does not include cadherin genes, but it does express C-type lectins (carbohydrate-binding proteins performing various functions in animals, including intercellular interaction, immune response and apoptosis). A reverse pattern of gene distribution is seen in *Pigoraptor*, where cadherin domain-containing transcripts were found but no C-type lectins (Hehenberger et al., 2017).

The presence of eukaryotrophy as a type of feeding within both filastereans and Pluriformea suggests that predation could be or have once been widespread among unicellular relatives of animals, and perhaps that the ancestor of Metazoa was able to feed on prey much larger than bacteria. The “joint feeding” we observed many times in cultures of *Syssomonas* or *Pigoraptor*, including the behaviour where cells are attracted to the large prey by other predators already feeding on it, is probably mediated by chemical signaling of the initially attached predator cell. The newly arriving cells also adhere to the plasmalemma of the prey, partially merging with each other and sucking out the contents of the large prey cell together. The merging of predator cells during feeding is quite unusual, and may represent a new factor to consider in the emergence of aggregated multicellularity. In addition, putative chemical signaling to attract other cells of its species is observed during the formation of syncytial structures in these species. In this context, alpha and beta-integrins and other components of the so-called integrin adhesome, which are responsible for interaction with the extracellular matrix and the transmission of various intercellular signals, were found in the transcriptomes of all three studied species (Hehenberger et al., 2017).

#### Starch breakdown by Syssomonas

An interesting phenomenon was observed in clonal cultures of *Syssomonas*, where the predator can completely engulf starch granules of the same size as the cell, also mediating the rapid destruction of rice grains into smaller fragments and individual starch crystals (Fig. S1). It is possible that *Syssomonas* secretes hydrolytic enzymes that provide near-membrane extracellular digestion. Near-membrane extracellular digestion in animals is of great importance for the breakdown of various biopolymers and organic molecules (for example, in the intestinal epithelium of mammals in the zone of limbus strigillatus in the glycocalix layer).

This ability of *Syssomonas* to feed on starch is likely promoted by the expression of numerous enzymes that are putatively involved in starch breakdown (several α-amylases and α-glucosidases, a glycogen debranching enzyme and a glycogen phosphorylase) (Table 1). For example, *Syssomonas* has five distinct putative alpha-amylases, one of them not found in any other Holozoa present in our database (Table 1). Similarly, one of the four α-glucosidases in *S. multiformis* seems to be specific to this lineage, and possibly the Filasterea, within the Holozoa. While α-amylases and α-glucosidases are able to hydrolyze α-1,4-linked glycosidic linkages, mobilization of the starch molecule at the α-1,6 glycosidic bonds at branch points requires the activity of debranching enzymes. A possible candidate for the catalysis of this reaction is a conserved glycogen debranching enzyme in *S. multiformis*, orthologous to the human *AGL* gene (Table 1). Additionally, we identified a transcript for a glycogen phosphorylase (orthologous to the human *PYGB, PYGL* and *PYGM* genes), an enzyme involved in the degradation of large branched glycan polymers.

**Table 1.** Transcripts for putative starch-degrading enzymes in *S. multiformis* and the presence/absence of corresponding orthologs in other unicellular holozoan lineages/Metazoa. Accession numbers for *S. multiformis* are from the corresponding TransDecoder output files as described in Hehenberger et al., 2017. Pfam domains used to identify candidates in *S. multiformis* and/or annotated direct human orthologous genes are indicated in brackets in column 1. NA, no direct orthologs recovered in our dataset; +, direct orthologs present in phylogeny; (+) short fragment of one lineage representative.

|  | *S. multiformis* | Ichthyosporea | Filasterea | Choanoflagellates | Metazoa |
| --- | --- | --- | --- | --- | --- |
| **α-amylases (PF00128)** |  |  |  |  |  |
| 1. | Colp12_sorted@267,<br>Colp12_sorted@268,<br>Colp12_sorted@269 | + | (+) | + | NA |
| 2. | Colp12_sorted@19603 | NA | NA | NA | NA |
| 3. | Colp12_sorted@18987,<br>Colp12_sorted@16467,<br>Colp12_sorted@17178 | NA | + | + | + |
| 4.= paralog to 3. | Colp12_sorted@8715 | + | + | + | + |
| 5. (*SLC3A1*) | Colp12_sorted@10150 | NA | NA | + | + |
| **α-glucosidases (PF01055)** |  |  |  |  |  |
| 1. | Colp12_sorted@9290,<br>Colp12_sorted@22295 | NA | (+) | NA | NA |
| 2. | Colp12_sorted@1339,<br>Colp12_sorted@1341 | + | + | NA | + |
| 3.= paralog to 2. | Colp12_sorted@9196,<br>Colp12_sorted@14409,<br>Colp12_sorted@18957,<br>Colp12_sorted@21457,<br>Colp12_sorted@31517 | + | + | NA | + |
| 4. (*GANAB/GANC*) | Colp12_sorted@13767 | + | + | + | + |
| **glycogen phosphorylase (PYGB/PYGL/PYGM)** |  |  |  |  |  |
| 1. | Colp12_sorted@1564 | + | + | + | + |
| **glycogen debranching enzyme (AGL)** |  |  |  |  |  |
| 1. | Colp12_sorted@14615 | + | + | + | + |

Our observations also show that, in the presence of starch in culture, *Syssomonas* can form resting stages of unidentified genesis, which tend to adhere to each other and to starch grains.

#### Structural features

*Syssomonas* and *Pigoraptor* both display a broad morphological plasticity: all three species have a flagellar stage, form pseudopodia and cysts, and can form aggregations of several cells. *Syssomonas multiformis* also has an amoeboid non-flagellar stage. The dominating life form of all three species in culture is the uniflagellar swimming cell. Interestingly, amoeboid and pseudopodial life forms were detected in cultures only after two years of cultivation and observation, suggesting they may be extremely rare in nature. Overall, the morphological differences between cells of the same type of the two genera, *Syssomonas* and *Pigoraptor*, are few and subtle. Given these genera are distantly related within the tree of Holozoa, it is interesting to speculate that they may be the result of morphostasis and by extension retain features resembling those of an ancestral state of holozoan lineages.

It has been established that single cells of *Syssomonas* and *Pigoraptor* can temporarily attach themselves to the substrate and, by beating their flagellum, can create water currents to putatively attract food particles, similar to choanoflagellates and sponge choanocytes (Fig. 1 E, Video 2). Choanocytes and choanoflagellates possess, in addition to the flagellum, a collar consisting of cytoplasmic outgrowths reinforced with actin filaments (microvilli) that serve to capture bacterial prey. The thin filopodia that are observed on the cell surface of all three *Syssomonas* and *Pigoraptor* species may thus be homologous to collar microvilli. But this will require further evidence in form of homologous proteins in these structures or evidence of their function in *Syssomonas* and *Pigoraptor*. While the filopodia of *Syssomonas* have no obvious structural contents, the outgrowths of *Pigoraptor* sometimes contain microtubular-like profiles. Cross sections of these structures were not obtained, but they may represent parallel microfilaments such as recently found in the filopodial arms of *Ministeria vibrans* (Mylnikov et al., 2019). The organisation of the *Ministeria* filopodial arms in turn resembles the microvilli of choanoflagellates, which have stable bundles of microfilaments at their base. It has been proposed previously that the ancestor of Filozoa (Filasterea+Choanoflagellida+Metazoa) probably had already developed filose tentacles, which have aggregated into a collar in the common ancestor of choanoflagellates/sponges (Shalchian-Tabrizi et al., 2008), and that microvilli were present in the common ancestor of Filozoa (Mylnikov et al., 2019).

A single, posterior flagellum is the defining characteristic of opisthokonts (Cavalier-Smith, Chao, 2003). However, the flagellum has not yet been found in all known Opisthokonta lineages. Torruella et al. (2015) have found several proteins corresponding to key components of the flagellum in *Corallochytrium* and the filose amoeba *Ministeria vibrans*, which have been considered to lack flagella. The authors have shown that the stalk used by *Ministeria* to attach to the substrate is a modified flagellum. Recently, morphological observations on another strain of *Ministeria vibrans* (strain L27; Mylnikov et al., 2019) revealed that this strain lacks the stalk for substrate attachment, but possesses a typical flagellum that projects forward and beats at attached to the substrate cells (see Fig. 2h and Video S1 in Mylnikov et al., 2019). The authors concluded that the filasterean ancestor possessed a flagellum, which was subsequently lost in *Capsaspora owczarzaki*. In the case of *Corallochytrium limacisporum*, it was suggested that it has a cryptic flagellate stage in its life cycle (Torruella et al., 2015), as has been proposed for other eukaryotes (*Aureococcus* and *Ostreococcus*, for instance) based on their genome sequences (Wickstead, Gull, 2012). Therefore, the flagellate stage could have been the one morphological trait uniting *Corallochytrium* and *Syssomonas* within “Pluriformea”. Interestingly, the ancestor of ichthyosporeans probably also had a flagellum, which is preserved in the Dermocystida (at the stage of zoospores), but was again lost in the Ichthyophonida.

The central filament of the flagellum, which connects the central pair of microtubules with the transversal plate in *Syssomonas* and *Pigoraptor*, is also noteworthy, since this character was previously known only in the choanoflagellates and was considered a unique feature for this lineage. The cone-shaped elevation of the surface membrane around the base of the flagellum in *Syssomonas* was also thought typical for choanoflagellates.

#### Origins of multicellularity

As mentioned above, numerous theories about the origin of Metazoa exist. One of the first and widely accepted evolutionary theories on the origin of animals is the Gastrea theory of Ernst Haeckel (Haeckel, 1874). Based on the blastula and gastrula stages that various animals undergo during their embryonic development, Haeckel suggested that the first step in the evolution of multicellularity in animals was the formation of a hollow ball, the walls of which consisted of identical flagellated cells, which he called Blastea. This stage is followed by gastrulation, where the ball invaginated, leading to the primary cellular differentiation into ecto- and endoderm. In combination with modern theories, such as the choanoblastea theory, which highlights the similarity between Haeckel’s Blastea and the choanoflagellate colony (Nielsen, 2008), this model is still the most commonly used explanation for the origin of multicellular animals. An important assumption of the Gastrea theory is that cell differentiation took place only *after* multicellularity arose, suggesting that animals originated from a single cell type. This in turn generated the hypothesis that multicellular animals originated from a choanoflagellate-like colony-forming ancestor (Sebé-Pedrós et al., 2017), supported by the possible homology between sponge choanocytes and choanoflagellates (Adamska, 2016), and is consistent with the idea behind the Haeckel-Muller Biogenetic Law, that ontogenesis recapitulates phylogenesis (Hashimshony et al., 2015).

However, there are, for example, some basic differences between sponges and choanoflagellates in how their collar and flagella interact, so, though choanocytes and choanoflagellates are superficially similar, homology should not be automatically assumed (see Mah et al., 2014 for details). More importantly, recent ultrastructural studies on the structure of kinetids (flagellar apparatus) of various sponge choanocytes and choanoflagellates show that they are fundamentally different in many respects (see Pozdnyakov, Karpov, 2016; Pozdnyakov et al., 2017, 2018 for details). These differences are significant, as the kinetid (consisting of the flagella itself, the transition zone, and the kinetosomes with attached microtubular or fibrillar roots) represents one of the very few ultrastructural systems in eukaryotic cells considered a conservative indicator of phylogenetic relationship (Lynn, Small, 1981; Moestrup, 2000; Yubuki and Leander, 2013). Choanocyte kinetids contain more elements that can be considered plesiomorphic for opisthokonts than do choanoflagellate kinetids. For example, significant differences in the spatial arrangement of the non-flagellar basal body, in the structure of the transitional zone and the flagellar root system were found when comparing the flagellate apparatus of the freshwater sponge *Ephydatia fluviatilis* (order Haplosclerida) and the choanoflagellates, but in all of these features the *Ephydatia* choanocyte kinetid is similar to the kinetid of zoospores of chytrids, which are more distantly-related Holomycota (Karpov, Efremova, 1994). Flagellated cells of some ichtyosporeans also possess ultrastructural features in common with flagellated fungi from Holomycota. Therefore, the idea that sponges and by extension all Metazoa descend directly from a single-celled organism similar to choanoflagellates is not supported by the results of the kinetid ultrastructure study (Pozdnyakov et al., 2017).

A more detailed understanding of the unicellular relatives of animals has, however, raised an alternative to the Gastrea theory. Specifically, the presence of diverse life forms in complex life cycles and the prevalence of cellular aggregations that bear little similarity to the blastula all suggest that cell *differentiation might have preceded* the origin of the blastula. Moreover, unicellular relatives of animals were shown to contain a variety of genes homologous to those involved in cell adhesion, differentiation and development, and signal transduction in Metazoa. Some of them were considered to be unique to animals (e.g. transcription factors T-box and Rel/NF-kappa B, Crumbs protein, integrin beta) as they were absent in choanoflagellates (the closest relatives of Metazoa), but later they were found to be present in other unicellular Holozoa (Mikhailov et al., 2009; Sebé-Pedrós et al., 2013a; Shalchian-Tabrizi et al., 2008). To date, it is well known that homologues of most genes controlling the development of animals, their cell differentiation, cell-cell and cell-matrix adhesion are present in various lineages of unicellular organisms (King et al., 2003; Sebé-Pedrós et al., 2016; Suga et al., 2013; Williams et al., 2014), which suggests that genetic programs of cellular differentiation and adhesion arose relatively early in the evolution of opisthokonts and before the emergence of multicellularity (King et al., 2003; Ruiz-Trillo et al., 2007; Shalchian-Tabrizi et al., 2008; Mikhailov et al., 2009; Brunet, King, 2018).

One specific idea positing that cell differentiation preceded the formation of colonies is the “synzoospore hypothesis” (Sachwatkin, 1956; Zakhvatkin, 1949; and see Mikhailov et al. 2009 for details). In brief, three types of cell cycle alternate in the ontogenesis of multicellular animals: monotomy (alternate phases of cell growth and division of somatic cells), hypertrophic growth (in female sex cells) and palintomy (the egg undergoes a series of consecutive divisions). Zakhvatkin noted that some protists alternate between different types of life cycle, and suggested that the unicellular ancestor of Metazoa already had differentiated cells as a result of such a complex life cycle. The life cycle complexity, in turn, results from the fact that monotomic cells are usually sedentary, or at least less mobile, and can change their phenotype (from flagellated to amoeboid, etc.) depending on the environment; the process of palintomy is necessary for the formation of morphologically identical dispersal cells (spores or zoospores). These dispersal cells remain attached to each other, forming a primary flagellated larva — the synzoospore or blastula (cited from Mikhailov et al., 2009).

The synzoospore hypothesis is consistent with recent observations of complex life cycles in unicellular opisthokonts possessing cellular differentiation, the presence of sedentary trophic phases, and a tendency to aggregation; as seen in choanoflagellates (Cavalier-Smith, 2017; Dayel et al., 2011; Dayel, King, 2014; Leadbeater, 1983; Maldonado, 2004), filastereans (Sebé-Pedrós et al., 2013b), *Corallochytrium* (Raghukumar, 1987), ichthyosporeans (Arkush et al., 2003; Ruiz-Trillo et al., 2007; Suga, Ruiz-Trillo, 2013), chytridiomycetes (Money, 2016), and nucleariid amoebae (Smirnov, 2000). According to this theory, multicellularity in animals arose through the temporal integration of various types of cells, which were already present in different parts of the life cycle. The hypothetical ancestor of animals in this model would thus already have genetic programs for cell differentiation (including cadherins, integrins, tyrosine kinases).

Developing the synzoospore hypothesis further, Mikhailov et al. (2009) proposed an evolutionary mechanism of “*transition from temporal to spatial cell differentiation*” to explain the emergence of multicellular animals. In this model, the ancestor of Metazoa was a sedentary colonial protist filter-feeder with colonies formed by cells of different types, which arose because filtration efficiency is significantly enhanced through the cooperation of cells of different types. Dispersal cells produced by the sedentary stage, the zoospores, remained attached together in early metazoans (which increased survivability) as a synzoospore to form a primary larva, the blastula. Development of a whole colony from such a multicellular larva occurred through the *differentiation of genetically identical* zoospore cells. This was critical for the maintenance of long-term cell adhesion and thus emergence of true multicellularity, as opposite to temporary colonies and aggregations composed of genetically heterogeneous cells. The authors suggest that the dispersal stages of the sedentary trophic body — primary blastula-like larvae – acquired adaptations to the predatory lifestyle, which triggered the development of primary intestine, muscular and nervous systems (Mikhailov et al., 2009).

The origin of multicellularity can in this view be seen as a *transition from temporal to spatiotemporal cell differentiation* (Sebé-Pedrós et al., 2017). In a unicellular ancestor of Metazoa that was a sexually reproducing bacteriotroph with many differentiated, temporally-separated cells, the transitions between different cell states would be regulated by expression of transcription factor families in response to environmental conditions such as availability of nutrients or preferred bacterial food. These temporally-regulated cell types then became spatially integrated, existing simultaneously but in different parts of a now multicellular conglomerate with different cell types carrying out different functions. Further diversification could then be accompanied by the evolution of additional mechanisms for complex gene regulation networks involving signaling pathways, expansion of transcription factors, and the evolution of new genomic regulatory mechanisms to control spatial differentiation of existing genetic modules specific to a particular cell type. At that point, the life cycle of the protozoan ancestor of animals probably included one or more clonal and/or aggregative multicellular stages (Sebé-Pedrós et al., 2017).

To distinguish between the models for the origin of animal multicellularity, genomics alone is not sufficient, and data on morphology, life cycle, and structural features of basal holozoans is also needed. From the current analysis, all three novel species of unicellular Holozoa have life histories that are consistent with major elements of the synzoospore model (see Fig. 3A in Mikhailov et al., 2009 and Fig. 5a,b in Sebé-Pedrós et al., 2017). Specifically, these organisms have complex life histories characterized by a variety of forms: flagellates, amoebae, amoeboflagellates, cysts. All three species have the tendency to form aggregations. *Syssomonas* possesses both clonal and aggregative multicellular stages, as predicted for the ancestor of animals. Moreover, the formation of aggregations can be associated with feeding on large eukaryotic prey, but also by cysts, which can adhere to each other and to starch grains in culture and divide multiply. Both these probably require cellular signaling. Eating a large eukaryotic prey, sometimes exceeding the size of a predator, also leads to hypertrophic cell growth (described as proliferative stage in Sebé-Pedrós et al., 2017) with a subsequent phase of palintomic division (in *Syssomonas*). All these characters are predicted by the synzoospore model. Interestingly, many of these are apparently triggered by the behaviour of feeding on large eukaryotic prey, highlighting this as is an interesting and potentially powerful trigger in general for the formation and development of aggregates (e.g., joint feeding) and clonal multicellularity (e.g., hypertrophic growth followed by palintomy), perhaps playing a role in the origin of multicellularity in ancestors of Metazoa.

### Concluding remarks

As we acquire more information about the biology of known unicellular relatives of animals, and even more importantly, describe diverse new species of unicellular Holozoa, more reliable models for the evolutionary histories of specific characteristics that contributed to the emergence of multicellularity in animals are possible. *Syssomonas* and *Pigoraptor* are characterized by complex life cycles, the formation of multicellular aggregations, and an unusual diet for single-celled opisthokonts (partial cell fusion and joint sucking of large eukaryotic prey), all of these features providing new insights into the origin of multicellularity in Metazoa.

Genomic and transcriptome analysis of unicellular relatives of animals have shown that genes encoding proteins for cellular signaling and adhesion, as well as genes for embryonic development of multicellular organisms, arose before the emergence of multicellular animals (King et al., 2003; Ruiz-Trillo et al., 2007; Shalchian-Tabrizi et al., 2008; Hehenberger et al., 2017). While these genes almost certainly have slightly different functions in protists than in animals, they nevertheless probably relate to the ability to recognize the cells of their own species, prey, or organic molecules and contribute to the formation of multicellular aggregations, thus increasing the organism’s ability to adapt to environmental change. As we learn more about the natural history and behaviour of these organisms, the importance of these processes becomes even more clear. The ancestor of Metazoa probably formed cells of various types that could aggregate and had molecular mechanisms of cell differentiation and adhesion related to those processes. Therefore, cellular differentiation likely arose before the emergence of multicellularity.

The feeding modes of the ancestral metazoan may also have been more complex than previously thought, including not only bacterial prey, but also larger eukaryotic cells and organic structures. Indeed, the ability to feed on large eukaryotic prey could have been a powerful trigger in the formation and development both aggregative and clonal multicellular stages that played important roles in the emergence of multicellularity in animals. Lastly, we wish to point out that other new and deep lineages of opisthokonts undoubtedly exist that have not yet been described, and each of these will play an important role in the development of hypotheses on the origin of multicellular animals in future.

## Materials and Methods

Novel unicellular opisthokont predators were found in freshwater biotopes in Vietnam and Chile. *Syssomonas multiformis* (clone Colp-12) was obtained from the sample of freshwater pool (11°23’08.0”N, 107°21’44.9”E; T = 39°C; pH = 7.18; DO (ppm) = 0.64; conductivity (µS/cm) = 281; TDS (ppm) = 140), Tà Lài, Cát Tiên National Park, Dong Nai Province, S.R. Vietnam on April 29, 2013. *Pigoraptor vietnamica* (clone Opistho-1) was obtained from freshwater Lake Dak Minh, silty sand on the littoral (12°54′50″N, 107°48′26″E; T = 27 °C; pH=7.03; DO (ppm) = 7.43; conductivity (µS/cm) = 109; TDS (ppm) = 54), Dak Lak Province, S.R. Vietnam on March 26 2015. *Pigoraptor chileana* (clone Opistho-2) was obtained from the bottom sediments of freshwater temporary water body (submerged meadow, 54°02’29.7”S, 68°55’18.3”W; T = 16.5°C; pH = 6.62; conductivity (µS/cm) = 141; TDS (ppm) = 72) near the Lake Lago Blanca, Tierra del Fuego, Chile on November 4, 2015.

The samples were examined on the third, sixth and ninth days of incubation in accordance with methods described previously (Tikhonenkov et al., 2008). Following isolation by glass micropipette, freshwater clones Colp-12, Opistho-1, and Opistho-2 were propagated on the bodonid *Parabodo caudatus* (strain BAS-1, IBIW RAS) grown in Pratt’s medium or spring water (Aqua Minerale, PepsiCo, Moscow Region, Russia or PC Natural Spring Water, President’s Choice, Toronto, Canada) by using the bacterium *Pseudomonas fluorescens* as food (Tikhonenkov et al., 2014). The clone Colp-12 was perished after five years of cultivation. The clones Opistho-1 and Opistho-2 are stored in the ―Live culture collection of free-living amoebae, heterotrophic flagellates and heliozoans” at the Institute for Biology of Inland Waters, Russian Academy of Science.

Light microscopy observations were made by using the Zeiss Axio Scope A.1 equipped with a DIC contrast water immersion objective (63x). The images were taken with the AVT HORN MC-1009/S analog video camera and directly digitized by using the Behold TV 409 FM tuner. Cells with engulfed starch granules were inspected by epifluorescence microscopy after DAPI staining using the Zeiss Axioplan 2 Imaging microscope.

For transmission electron microscopy (TEM), cells were centrifuged, fixed at 1 °C for 15-60 min in a cocktail of 0.6% glutaraldehyde and 2% OsO_4_ (final concentration) prepared using a 0.1 M cacodylate buffer (pH 7.2). Fixed cells were dehydrated in alcohol and acetone series (30, 50, 70, 96, and 100%, 20 minutes in each step). Afterward, the cells were embedded in a mixture of Araldite and Epon (Luft, 1961). Ultrathin sections were prepared with an LKB ultramicrotome (Sweden) and observed by using the JEM 1011 transmission electron microscope (JEOL, Japan).

For scanning electron microscopy (SEM), cells from exponential growth phase were fixed as for TEM but only for 10 min at 22 °C and gently drawn onto a polycarbonate filter (diameter 24 mm, pores 0.8 µm). Following the filtration, the specimens were taken through a graded ethanol dehydration and acetone, and finally put into a chamber of a critical point device for drying. Then dry filters with fixed specimens were mounted on aluminum stubs, coated with gold-palladium, and observed with a JSM-6510LV scanning electron microscope (JEOL, Japan).

Analysis of enzymes involved in starch breakdown was based on transcriptomic data obtained as described earlier (Hehenberger et al., 2017). To identify candidates putatively involved in starch breakdown, we used the results of a previous hmmscan analysis of *S. multiformis* (Hehenberger et al., 2017) to search for Pfam domains present in known starch-degrading enzymes/enzyme families, such as α-amylases (PF00128), glycoside hydrolase families containing α-glucosidases (PF02056, PF01055, PF03200, PF10566), α-glucan water dikinase 1 (*GWD1*, PF01326), phosphoglucan phosphatase (*DSP4*, PF00782 and PF16561), disproportionating enzymes (PF02446) and pullulanases (PF17967). Additionally, we submitted the *S. multiformis* sorted transcriptome to the KEGG Automatic Annotation Server (KAAS) (Kanehisa et al., 2014) for functional annotation and investigated the output for transcripts involved in starch metabolism. All candidates were investigated using phylogenetic reconstruction. Briefly, they were used as queries in a BLASTp search (e-value threshold 1e-5) against a comprehensive custom database containing representatives of all major eukaryotic groups and RefSeq data from all bacterial phyla at NCBI (https://www.ncbi.nlm.nih.gov/, last accessed December 2017) (Altschul et al., 1990). The database was subjected to CD-HIT with a similarity threshold of 85% to reduce redundant sequences and paralogs (Li and Godzik, 2006). Results from blast searches were parsed for hits with a minimum query coverage of 50% and e-values of less than 1e-5. The number of bacterial hits was restrained to 20 hits per phylum (for FCB group, most classes of Proteobacteria, PVC group, Spirochaetes, Actinobacteria, Cyanobacteria (unranked) and Firmicutes) or 10 per phylum (remaining bacterial phyla) as defined by NCBI taxonomy. Parsed hits were aligned with MAFFT v. 7.212, using the–auto option, poorly aligned regions were eliminated using trimAl v.1.2 with a gap threshold of 80% (Katoh and Standley, 2013; Capella-Gutiérrez et al., 2009). Maximum likelihood tree reconstructions were then performed with FastTree v. 2.1.7 using the default options (Price et al., 2010). Phylogenies with overlapping taxa were consolidated by combining the parsed hits of the corresponding queries, removing duplicates and repeating the alignment, trimming and tree reconstruction steps as described above.

## Acknowledgements

We thank Kristina I. Prokina for sample collection in Chile, Dr. Hoan Q. Tran and Tran Duc Dien for assistance with sample collection and trip management in Vietnam, Jürgen F.H. Strassert for help with DAPI staining, and Vladimir V. Aleshin and Kirill V. Mikhailov for fruitful discussion on different aspects of origin of Metazoa. Field work in Vietnam is part of the project “Ecolan 3.2” of the Russian-Vietnam Tropical Centre.

## Legends for videos

Video 1. Swimming of *Syssomonas multiformis* cell with rotation.

Video 2. Attached cell of *Syssomonas multiformis* and rapid flagellum beating.

Video 3. Amoeboflagellate stage of *Syssomonas multiformis.* Cells of eukaryotic prey *Parabodo caudatus* are also visible.

Video 4. Loss of flagellum in *Syssomonas multiformis* and transition to amoeba.

Video 5. Transformation of amoeba into a cyst in *Syssomonas multiformis*.

Video 6. Palintomic divisions inside the cyst of *Syssomonas multiformis*.

Video 7. Division into two cell structures in *Syssomonas multiformis.*

Video 8. Cell and cyst of *Syssomonas multiformis* with vesicular structures inside.

Video 9. Feeding of *Syssomonas multiformis* on eukaryotic prey.

Video 10. Feeding of *Syssomonas multiformis* on bacteria.

Video 11. Temporary cell aggregations of *Syssomonas multiformis.*

Video 12. Floating rosette-like aggregation of *Syssomonas multiformis.*

Video 13. Syncytium-like structures and budding of young flagellated daughter cells in *Syssomonas multiformis.*

Video 14. Joint feeding of *Pigoraptor vietnamica* on died cell of *Parabodo caudatus.*

Video 15. Joint feeding of *Pigoraptor chileana* on died cell of *Parabodo caudatus.*

Video 16. Temporary cell aggregation of *Pigoraptor chileana.*

***Videos are available at* https://drive.google.com/drive/folders/1ewNvwSQCAVG81eYjJPdhR41aBpUKR05L**

